# CRZ1 regulator and calcium cooperatively modulate holocellulases gene expression in *Trichoderma reesei* QM6a

**DOI:** 10.1101/622647

**Authors:** Leonardo Martins-Santana, Renato Graciano de Paula, Adriano Gomes Silva, Douglas Christian Borges Lopes, Roberto do Nascimento Silva, Rafael Silva-Rocha

**Affiliations:** Department of Biochemistry and Immunology, University of São Paulo, Medical School of Ribeirão Preto, São Paulo, Brazil; Department of Cell and Molecular Biology and Pathogenic Agents, Systems and Synthetic Biology Laboratory, University of São Paulo, Medical School of Ribeirão Preto, São Paulo, Brazil

**Keywords:** Gene regulatory network, cellulase expression, transcriptional regulation, calcium sensing

## Abstract

**Background:** *Trichoderma reesei* is the main filamentous fungus used in industry to produce cellulases. Over the last decades, there have been a strong increase in the understanding of the regulatory network controlling cellulase-coding gene expression in response to a number of inducers and environmental signals. In this sense, the role of calcium and the Calcineurin-responsive protein (CRZ1) has been investigated in industrially relevant strains of *T. reesei* RUT-C30, but this system has not been investigated in wild-type reference strain of this fungus.

**Results:** Here, we investigated the role of CRZ1 and Ca^2+^ signaling in the fungus *T. reesei* QM6a. For this, we first searched for potential CRZ1 binding sites in promoter regions of key genes coding holocellulases, as well as transcriptional regulators and sugar and calcium transporters. Using a nearly constructed *T. reesei* Δ*crz1* strain, we demonstrated that most of the genes expected to be regulated by CRZ1 were affected in the mutant strain induced with sugarcane bagasse (SCB) and cellulose. In particular, our data demonstrate that Ca^2+^ acts synergistically with CRZ1 to modulate gene expression, but also exerts CRZ1-independent regulatory role in gene expression in *T. reesei*, suggesting the existence of additional Ca^2+^ sensing mechanisms in this fungus.

**Conclusions:** This work presents new evidence on the regulatory role of CRZ1 and Ca^2+^ sensing in the regulation of cellulolytic enzymes in *T. reesei*, evidencing significant and previously unknown function of this Ca^2+^ sensing system in the control key transcriptional regulators (XYR1 and CRE1) and on the expression of genes related to sugar and Ca^2+^ transport. Taken together, the data obtained here provide new evidence on the regulatory network of *T. reesei* related to plant biomass deconstruction.

## Background

Cellulases and hemicellulases are key enzymes for the production of second generation ethanol production worldwide [1]. The promising utilization of agroindustrial residues as energy source has triggered a tremendous interest for engineering microorganisms for the low cost production of these enzymes aiming the conversion of complex plant biomass into fermentable sugars, which in turn could be used for ethanol production [2–4]. In this sense, the saprophyte organism *T. reesei* is one of the the main filamentous fungus used in industry to produce these enzymes [2]. Despite the great potential of this fungus for high performance cellulase and hemicellulase production, the regulatory network controlling this system is a relevant field of study and not fully understood yet. Additionally, the comprehension of transcriptional networks which coordinate the transcription of holocellulases is one of the main focus to allow enzyme production in a viable at industrial scale [5].

*T. reesei* possess a great capability for enzyme secretion [6] but few cellulases and hemicellulases genes. Even harboring this limited gene repertoire, its regulation is highly precise and controlled at molecular levels by external signals [7]. Several transcription factors (TFs) are now known for exert modulation on the expression levels of holocellulases genes [8]. In this sense, the regulatory network which encompasses cell wall deconstruction is under regulation of positive regulators, as XYR1 [9], ACEII [10], LAE1 [11], BglR [12], VEL1 [13], HAP 2/3/5 complex [14] and other proteins which play a negative control in such process, as CRE1 [15], ACEI [16] and RCE1[17]. Additionally, it is already known that xylanases genes may also be exclusive targets of regulation byTF such as Xpp1[18] and SxlR [19]. Interestingly, the same regulators are not seem to be responsible for influencing cellulases expression in the same fungus.

The complex network that coordinates holocellulases expression in *T. reesei* is also under regulation of nutritional variability. Therefore, differential gene expression was discussed by Castro *et al.* [20] and Antonieto *et al.* [21] in studies which reported the influence of carbon source on holocellulases transcription. Robust techniques as RNA sequencing have already highlighted the importance of sugar availability on the regulation mediated by transcription factors in *T. reesei* [22], evidencing the growing need of elucidating deep layers of regulation in this fungus.

Nutritional requirements and pH stress conditions control are target of research among a wide number of fungal species, such as *Neurospora crassa* [23], *Trichoderma harzianum* [24] and *T. reesei* [25]. However, the influence of several stress conditions, such as the effect of ions concentration exposure in transcription, still remains poorly understood in the industrial workhorse *T. reesei*. Despite not completely elucidated, recent studies have demonstrated the balance mediated by signaling pathways to keep physiological homeostasis when *T. reesei* is exposed to Ca^2+^ stress through Calmodulin-Calcineurin downstream activation steps [26].

CRZ1 (Calcineurin responsive protein) is a key target of Calcineurin (CN) in fungal cells and is a well-conserved orthologue protein in several fungi species [27–29], being responsible for many coordinated cellular functions. In *Saccharomyces cerevisiae*, a deletion of this protein resulted in hypersensitivity to chloride and chitosan, as well as mating and transcriptional response to alkaline and stress conditions defects [30]. In another *crz1* mutant yeast, *Candida albicans*, the knockout effect was correlated to changes on the pattern of sensitiveness to Ca^2+^, Mn^2+^ and Li^2+^, as well as in hypersensitiveness to azoles molecules [31]. The role of CRZ1 in filamentous fungi are already described for some species, and in this context, for the pathogenic and opportunistic *Aspergillus fumigatus, crz1* knockout resulted in virulence attenuation and developmental defects in both hyphal morphology and conidiation [32]. Effects of CRZ1 in filamentous fungi were also mentioned for *Cryptococcus neoformans*, where growth at temperatures higher than 39°C was compromised in *crz1* knockout strains, as well as its absence resulted in sensitiveness to cell wall perturbing agents [29,33].

In *T. reesei*, the role of Ca^2+^-CaM/CN/CRZ1 pathway is poorly studied and established in comparison to other fungal species. Yet, it is already known that Ca^2+^ signaling pathway may exert effects on the modulation of holocellulases gene expression in the industrial strain of *T. reesei* RUT-C30 [26]. Therefore, Chen *et al.* [26] showed that the regulation of relevant industrial genes, such as *cbh1, eg1* and *xyr1* genes were negatively modulated in the absence of CRZ1 when calcium was added to a cellulose-supplemented medium.

Despite the evidence that CRZ1 and Ca^2+^ may improve holocellulases expression, these results were obtained when the industrial strain of *T. reesei* RUT-C30 was evaluated in different environmental conditions. Yet, this strain harbors a deletion of approximately 85 kilobases (kb) from *T. reesei’s* genome which includes the *cre1* locus, resulting in phenotypes that are independent on CRE1-mediated influence [34,35]. Regarding, the role of CRZ1 and Ca^2+^ have not been investigated in wild type strain of *T. reesei* (QM6a), which harbors the full set of regulatory components into its genome. In order to elucidate this, in our study, we investigate the role of CRZ1/Ca^2+^ in a wild set genes coding for transcriptional factors (TFs), enzyme and transporters related to holocellulose deconstruction. In this study, we were able to identify CRZ1 transcription factor binding sites (TFBS) in promoter regions of the differentially holocellulases expressed genes after *T. reesei* QM6a exposure to SCB and cellulose as carbon source. Additionally, we reported the fundamental role of CRZ1 on transcriptional regulation of such genes, as well as we demonstrated that Ca^2+^ can cooperatively with CRZ1, induces or represses the transcription of holocellulases genes. These findings bring evidences of Ca^2+^ to be believed as a pivotal ion in the control of stimuli concerning expression of sugar and calcium transporters in *T. reesei* QM6a cells in an CRZ1-independent but synergistic mechanism.

## Material and Methods

### Analysis of promoters sequence

For the identification of potential CRZ1-binding elements, sequences of 1kb upstream of the ATG from the Open Reading Frames (ORFs) of the promoter of TFs, hollocellulases and transporter genes as well as for homologue of calcium and sugar transporters were retrieved from the reference genome of *T. reesei*. Next, Position Weight Matrix (PWM) from CRZ1 of *S. cerevisiae* were obtained from YeTFaSCo (The Yeast Transcription Factor Specificity Compendium) collection [36] and used to calculate the scores in all promoters of *T. reesei*. Scores greater than zero were selected and normalized, revealing sequence with the higher probability of harboring a functional *cis*-regulatory element than a random motif. The normalization function was: Score_Normalized_(Xij) = Score(Xij)/MaxScore, being X the set of sense and antisense upstream regulatory sequences; *i* the sequence index; *j* the initial position for score calculation; and MaxScore the maximum score possible of the CRZ1 TF PWM. Through the histogram analyses of the normalized score, we established a threshold of 0.8, using a comparison between real sequences and a control dataset formed by random sequences. Finaly, tis threshold was used to map potetinal cis-regulatory elements for CRZ1 in the target of interest.

### Strains and media

*T. reesei* strain *QM6a*Δ*tmus53*Δ*pyr4* was used as wild type (WT) strain in this study and it was obtained from the Institute of Chemical Engineering (Viena University of Technology, Viena, Austria) [37]. The strain was maintained in MEX medium (3% (w/v) malt extract and 2% (w/v) agar)) at 30°C supplemented with 5 mM uridine for *pyr4* auxotrophic selection [38]. *T. reesei QM6a*Δ*tmus53*Δ*pyr4* wildtype and Δ*crz1* (constructed as described below) strains were grown on MEX medium at 30°C for 7-10 days until complete sporulation. For gene expression assays, approximately 10^6^ spores/mL from both strains were inoculated in Mandels Andreotti (MA) medium [39] containing 1% (w/v) sugarcane bagasse, cellulose (Avicel, Sigma-Aldrich) or glycerol as carbon source. For induction of gene expression in the presence of sugarcane bagasse, the substrate *in natura*, gently donated by Nardini Agroindustrial Ltd., Vista Alegre do Alto, São Paulo, Brazil, was previously prepared according to Souza et al. [40]. Mycelia from both strains (WT and Δ*crz1*) were pre-grown in MA medium supplemented with 1% (w/v) glycerol, in the presence and in the absence of 10 mM CaCl_2_, in orbital shaker at 30°C for 24 hours under 200 rpm. In these assays, MA medium presented a basal concentration of 2.6 mM Ca^2+^ to ensure the maintenance of fungal cellular viability and growth. The obtained mycelia were harvested and transferred to MA medium supplemented with 1% (w/v) sugarcane bagasse or cellulose (Avicel, Sigma-Aldrich) in orbital shaker at 200 rpm, 30°C for 8 h. All experimental conditions described were performed with three biological replicates. The resulting mycelia were collected by filtration, frozen in liquid nitrogen and stored at −80°C for RNA extraction procedures.

### Construction of crz1 deletion cassette and T. reesei genetic transformation

Deletion of *crz1* gene from *T. reesei QM6a*Δ*tmus53*Δ*pyr4* was performed as previously described [41]. For this, we design a deletion cassette containing the *pyr4* gene (orotidine-5′-phosphate decarboxylase gene of *T. reesei* - *Trire_74020*) as auxotrophic selection marker. All sequences of this cassette are available at the *T. reesei* genome database (https://genome.jgi.doe.gov/Trire2/Trire2.home.html) and the primers used in this study were designed to contain overlapping regions to the components sequence of the cassette (**Additional file 1: Table S1)**. The deletion cassette is composed by 1226 base pairs (bp) upstream to the 5’ *crz1* CDS flanking region (*Trire_36391*) and 1465 bp downstream to the *crz1* ORF 3’ flanking region. These two sequences flanks *pyr4* gene sequence, which consists of 1100 bp upstream and 1000 bp downstream to the *pyr4* CDS sequence in *T. reseei*’s genome. Individual fragments were amplified at 60°C with Phusion® High-Fidelity Polymerase (New England Biolabs) using *T. reesei QM6a*Δ*tmus53*Δ*pyr4* genomic DNA as template, according to manufacturer’s instructions. The PCR fragments were purified using QIAquick PCR Purification Kit (Qiagen). The assembly of the components was performed through homologous recombination in *S. cerevisiae*, as previously reported [42–44]. For this purpose, we used the pRS426 shuttle vector [45] previously treated with *Eco*RI and *XhoI* enzymes (New England Biolabs). These treatment resulted on the occurrence of cohesive ends correspondent to the external extremities in the primers used to obtain the 5’ and 3’ *pyr4* ORF flanker regions of the deletion cassette. Yeast transformation procedures were performed as described by Gietz and Schiestl, (2007) in the yeast *S. cerevisiae* strain SC9721 (*MATα his3-Δ200 URA3-52 leu2Δ1 lys2Δ202 trp1Δ63*) (Fungal Genetic Stock Center). The transformants were selected on YPD (1% (w/v) yeast extract, 2% (w/v) peptone and 1% (w/v) glucose) medium supplemented with lysine, histidine, leucine, and tryptophan and in the absence of uracil.

To confirm the complete assembly of the cassette, genomic DNA of *S. cerevisiae* was extracted according to Goldman et al. [46]. Primers 5’P*crz1* and 3’T*crz1* (**Additional file 1:Table S1)** were used to amplify the deletion cassette from the yeast genome using Phusion® High-Fidelity Polymerase (New England Biolabs) according to manufacturer’s instructions. The resultant amplicons were purified using QIAquick PCR Purification Kit (Qiagen) and were stored at −20°C until *T. reesei* transformation. Transformation of *T. reesei* was performed using 10 μg of the linearized deletion cassette by protoplast fusion methodology [38]. Transformants were selected in MA medium plates supplemented with 1% (w/v) glucose and 2% (w/v) agar (Sigma-Aldrich). Selection was carried out in the absence of uridine. Genomic DNA was extracted to confirm the cassette integration [46]. To verify the correct orientation of cassette positioning in the positive transformants, it was performed two PCR reactions with primers 5’P*crz1* Check1/3’*pyr4* Check 1 and 5’*pyr4* Check 2/3’T*crz1* Check 2 (**Additional file 1: Table S1, Additional File 2:Figure S1 A)**, which are specific for *pyr4* annealing regions and for external annealing regions into promoter and terminators sequences. A PCR to confirm the deletion of the *crz1* from *T. reesei* genome was performed using the primers 5’*crz1* ORF and 3’ *crz1* ORF (**Additional file 1: Table S1, Additional File 2:Figure S1 B)**. Null expression of *crz1* gene was also investigated with a quantitative PCR with total RNA using the primers PF *crz1* qRT-PCR and PR *crz1* qRT-PCR (**Additional file 1: Table S1, Additional File 2:Figure S1 C**).

### Gene expression analysis by Real-Time PCR

For gene expression analysis, 1 μg of total RNA of each culture condition was treated with DNAse I (Sigma-Aldrich). cDNA samples were synthesized using Maxima™ First Strand cDNA Synthesis kit (ThermoFisher Scientific) and posteriorly diluted 1:50 in DEPC water. For gene expression levels detection, SsoFast™ EvaGreen® Supermix (Bio-Rad) was used according to manufacturer’s instructions. A list of target genes whose expression were evaluated are available in **Additional file 3:Table S1**. Reactions were carried out at 95°C for 10 minutes, followed by 40 cycles at 95°C for 10 seconds and 60°C for 30 seconds in a Bio-Rad CFX96™ Real-Time System coupled to a C1000™ Thermal Cycler (BioRad). Expression of target genes was normalized by the β-actin endogenous transcript levels for each RNA prevenient from the culture conditions [47]. For sugarcane bagasse and cellulose expression quantification (for both in presence or in the absence of 10 mM calcium chloride) the 2^−ΔΔCt^ method was employed [48]. The quantification was relative to transcript levels observed in the WT and Δ*crz1* strains grown in glycerol for 24 h (for both in the presence or in the absence of 10 mM calcium chloride).

### Enzymatic activity assays

The activity of β-glucosidases and β-xylosidases were measured as the capacity of hydrolyzing pNPG and pNPX substrates as previously described [20,49,50]. Endoglucanase activity was determined using Carboxymethylcellulose (CMC) (Sigma-Aldrich) as substrate in microplates, in a protocol adapted from Xiao *et al.* [51]. We solubilize 1% (w/v) CMC in pH 4.8 sodium acetate buffer. After addition of 30 μL of culture supernatant, reactions were incubated for 30 minutes at 50°C. After this time, 60 μL of DNS were added to the reaction and the mixture were submitted to incubation for 5 minutes at 95°C [52]. Absorbance at 540 nm were used for samples measurements. One enzyme unit was defined as the amount of enzyme capable of releasing 1 μmol of reducing sugar per minute. Endoxylanases activity was determined using xylan from beechwood (Sigma-Aldrich) solubilized in pH 5.0 100 mM sodium acetate buffer as substrate. Reactions were performed with 25 μL of the culture supernatant and 50 μL of substrate at 50°C for 30 minutes. After incubation, we mixed 75 μL of DNS and heated the reactions at 95°C for 5 minutes to visualize the effects of reduction of dinitrosalicylic acid. Absorbances were carried at 540 nm. One unit of enzyme was defined as the amount of enzyme capable of releasing 1 μmol of reducing sugar per minute [53].

## Results

### Identification of crz1 binding motifs in T. reesei promoters

We first investigated the existence of putative binding sites for CRZ1 in *T. reesei* holocellulases and calcium and sugar transporter promoters using the defined sequence for *S. cerevisiae*. For this, we selected 30 genes related to biomass degradation in plant cell wall through proteins secreted by *T. reesei*, such as cellulases, hemicellulases and sugar transporters, as well as calcium transporters and TFs which play a significant role in this process. In this sense, we were able to identify potential *cis*-regulatory elements for CRZ1 in 24 of these sequences, as represented in Figure 1A-B. Interestingly, the major numbers of potential cis-regulatory elements for CRZ1 were observed in *cre1* and *cel7a* genes as well as for *Trire_56440* and *Trire_58952* calcium transporter proteins promoters (Figure 1C). We also observed that the promoter of *crz1* gene harbors 3 potential cis-regulatory elements in antisense orientation, suggesting that CRZ1 plays autoregulation at transcriptional level (Figure 1C). Taken together, these results evidenced the existence of putative binding sites for CRZ1 in several genes related to gene regulation, biomass and calcium transport, highlighting the potential role of this TF in the regulation of these genes.

**Figure 1.**
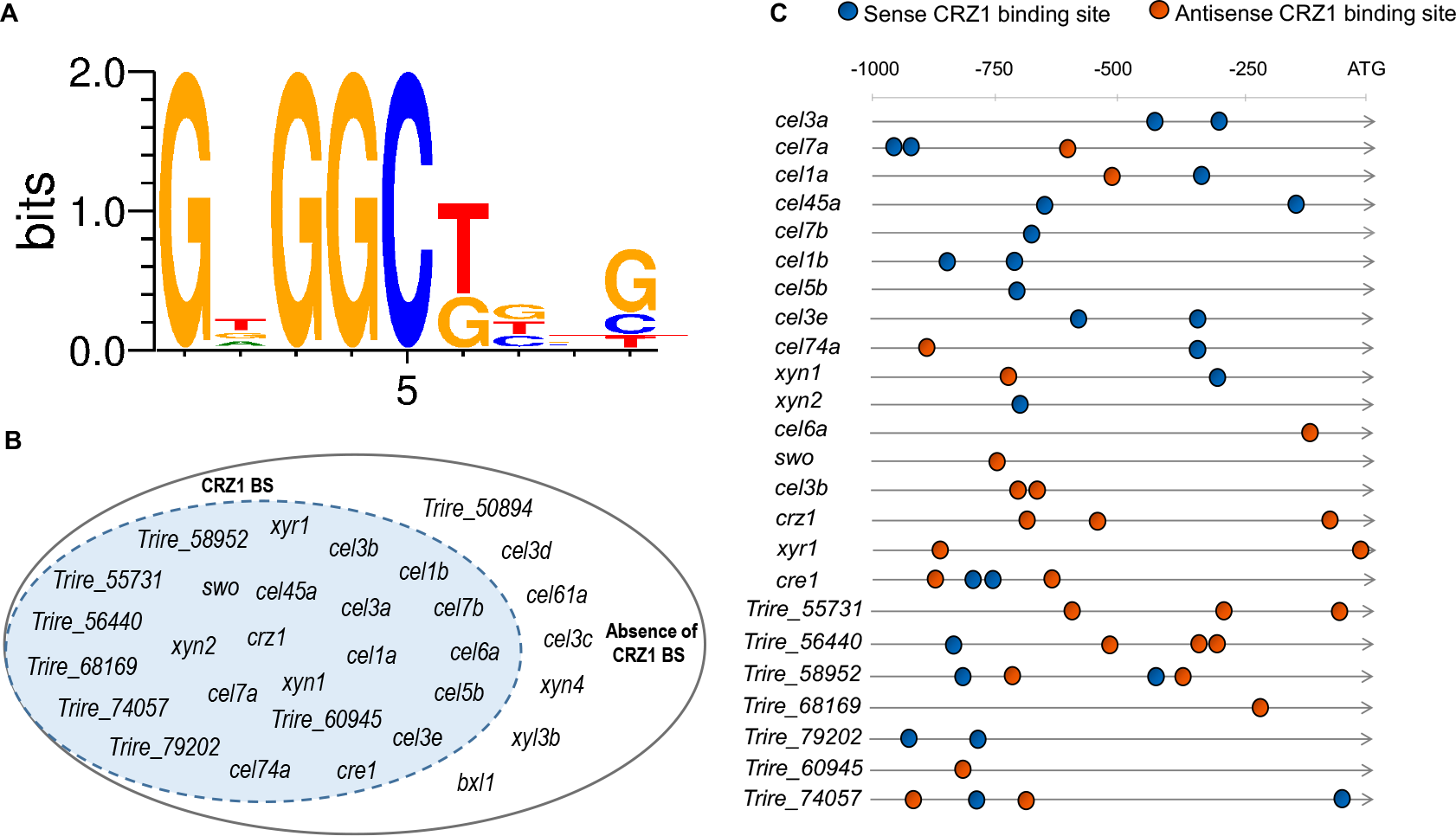
**A** – DNA Consensus sequence of CRZ1 binding in *T. reesei* obtained through bioinformatics analysis. **B** – Representative scheme of genes which harness CRZ1 BS (CRZ1 binding sites) at genomic level. **C** – *in silico* Identification of CRZ1 BS in promoter regions of holocellulases and calcium and sugar transporter genes in *T. reesei*. Blue spheres indicate that those sequences (previously shown in 1A) have been found in sense orientation and orange spheres represent the antisense sequences occurrence. The number of spheres represents the absolute frequency that these sequences were observed in bioinformatics analysis.

### crz1 and Ca^2+^ synergistically regulates TFs and cellobiohydrolase coding genes

Once we have identified putative binding sites for CRZ1 at promoter regions of the genes of interest, we tried to understand how carbon source, as well as Ca^2+^ supplementation, could modulate the transcription of these genes in the wild type and *Δcrz1* mutant background. In this sense, we observed that under exposure to SCB, the expression of *xyr1* was positively modulated in the wild type strain of *T. reesei* by the supplementation of 10 mM Ca^2+^ to the medium (Figure 2A). However, this stimulatory effect was lost in the *T. reesei Δcrz1* strain, indicating that this process was completely dependent on CRZ1. Additionally, the same effect was observed when Avicel was used as inducer (**Additional file 4: Figure S1 A**). In the case of *cre1*, we observed only minor changes in the expression of this TF under all conditions tested, with a small but significant reduction in *its* expression in the wild type strain upon Ca^2+^ induction or between wild type and *Δcrz1* strain without the exposure to this ion (Figure 2B). Yet, when its expression was assayed under Avicel exposure, no significant change was observed between any condition analyzed (**Additional file 4: Figure S1 B**). It is worth to notice that while putative CRZ1 binding sites were identified at both *xyr1* and *cre1* promoters (Figure 1C), only the former seems to be significantly regulated by this TF under the conditions analyzed here.

**Figure 2.**
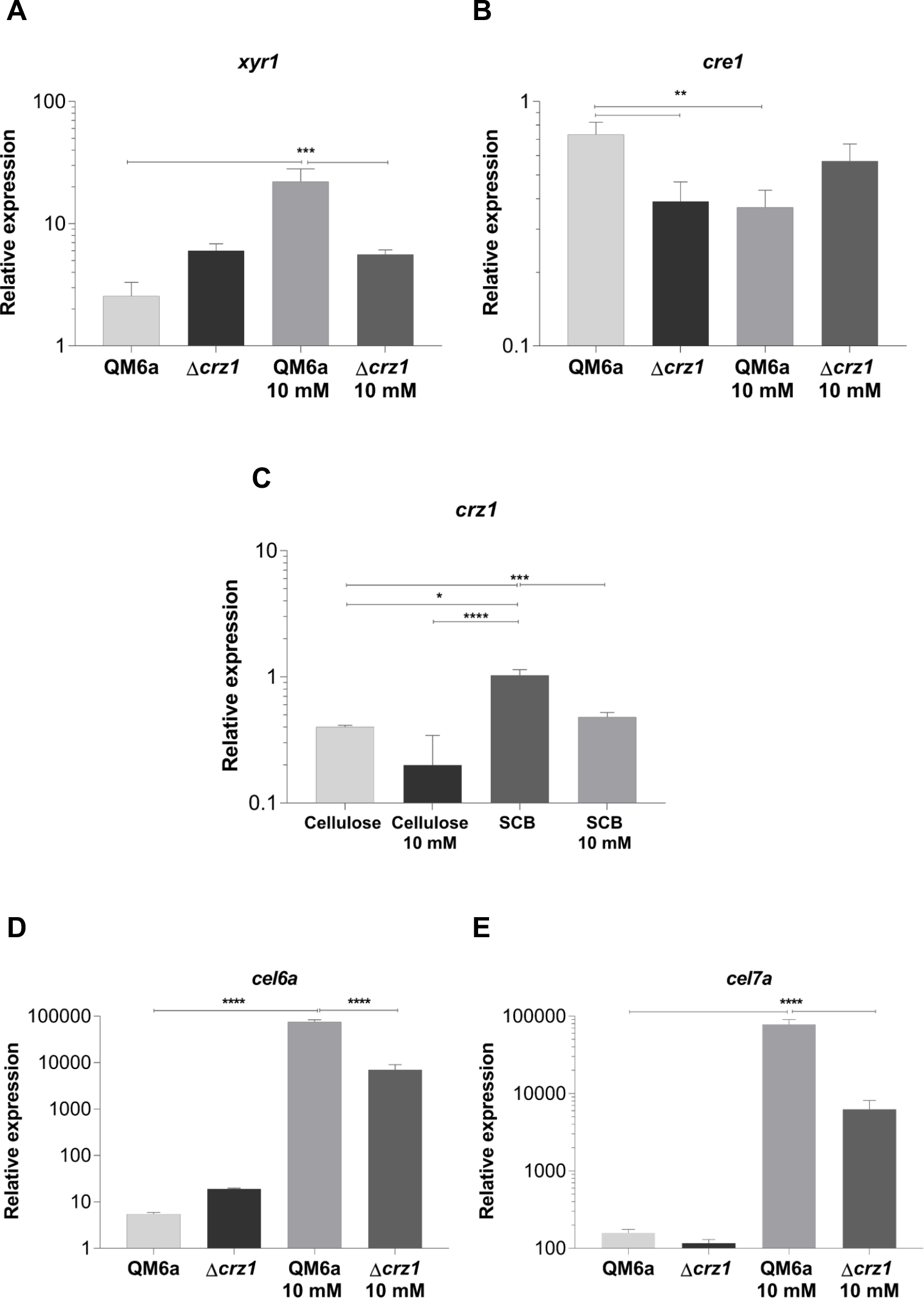
Analysis of differential expression of the transcription factors XYR1 (**A**), CRE1(**B**), and CRZ1 (**C**) of *T. reesei* and of cellobiohydrolases CEL6A (**D**) and CEL7a (**E**) from QM6a and *Δcrz1 T. reesei* strains after 8 h of growing in SCB and SCB supplemented with 10 mM Ca^2+^. CRZ1 expression in the presence of cellulose and the effect caused by calcium on this induction is also shown in panel C. Expression values are represented as log_10_ means of three biological replicates with standard deviation normalized by glycerol expression levels at the same condition. Statistical significance is represented as asterisks, considering p-value as at least < 0.05 (* < 0.05 < ** < 0.005 < *** < 0 0001 < ****)

Since we identified putative CRZ1 binding sites in *crz1* promoter, we were also interested in investigate how CRZ1-mediated autoregulation in *T. reesei* was related to carbon source availability. In this sense, Figure 2C shows the result only for wild type strain of *T. reesei* with or without Ca^2+^ exposure. As shown by the figure, under SCB induction, the expression levels of *crz1* were higher than those obtained after cellulose induction. In addition, after supplementation of the medium with 10 mM Ca^2+^, the expression of this gene decreased in the presence of SCB, suggesting that Ca^2+^ may exert a modulatory effect on CRZ1 autoregulation.

Since we observed that *xyr1* expression was induced by Ca^2+^ in a CRZ1 dependent manner, we next evaluated the role of these two components in the expression of *cel6a* and *cel7a*, the two major cellobiohydrolases from *T. reesei*. We found that Ca^2+^ supplementation resulted in a significant increase in *cel6a* and *cel7a* induction both under exposure to SCB (Figure 2D-E) and Avicel (**Additional file 4: Figure S1 C-D**). Additionally, deletion of crz1 resulted in a significant reduction in this stimulatory effect, even though this was not completely abolished. Taken together, these results indicated that CRZ1 and Ca^2+^ synergistically control the expression of celobiohydrolases coding genes under SCB and Avicel exposure, and highlight that an additional Ca^2+^-dependent and CRZ1-independent mechanism could take part in this process.

### Other cellulases genes are also key of regulation by crz1 and Ca^2+^

Since we observed that CRZ1 is capable to modulate *xyr1* expression levels, we next investigated how the expression of xylanase coding genes was modulated as a result of such effect mediated by CRZ1. Thus, under exposure to SCB, we observed a significant increase on the expression of these genes induced by the simple supplementation with Ca^2+^ (Figure 3A-E). Additionally, the deletion of *crz1* was not able to result in differences on the expression of these genes, but we detected a loss in their expression in the *T. reesei Δcrz1* strain after Ca^2+^ supplementation. Taking in advantage of these results, our findings suggest that the regulation of xylanase genes under SCB exposure is modulated by Ca^2+^ in a synergistically CRZ1-dependent mechanism, highlighting the relevance of this TF on a complete background concerning gene regulation in the wild type strain of this fungus. Interestingly, no effects were observed when Avicel was used as inducer, since none differential expression was detected under cellulose exposure (**Additional file 5: Figure S1 A-D**).

**Figure 3.**
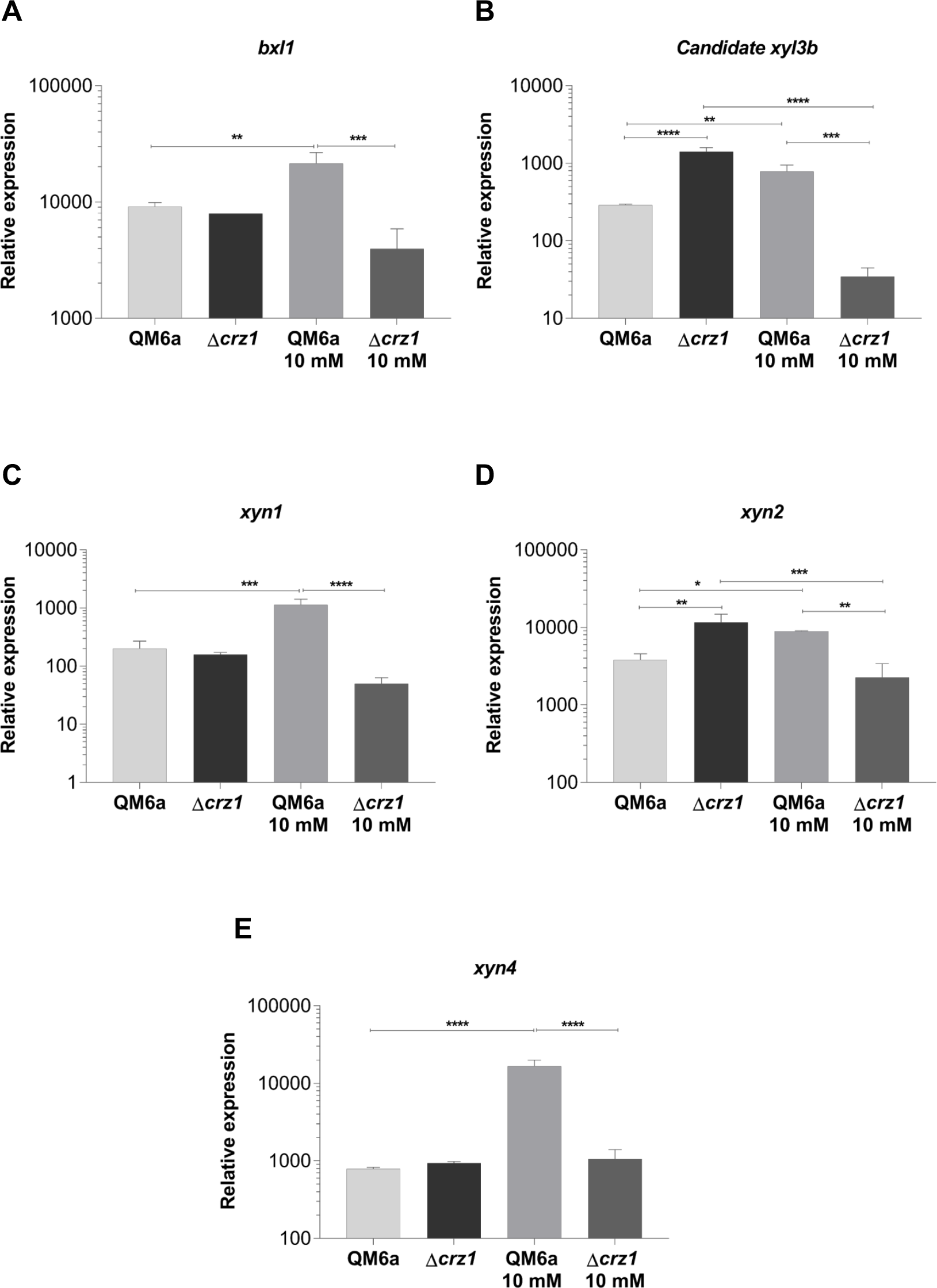
Relative expression of the β-xylosidase 1 and (**A**) the candidate *xyl3b* xylosidase (**B**) and of the endoxylanases *xynl* (**C**), *xyn2* (**D**) and *xyn4* (**E**) genes from QM6a and *Δcrzl T. reesei* strains after 8 h of induction SCB and SCB supplemented with 10 mM Ca^2+^. Expression values are represented as log_10_ means of three biological replicates with standard deviation normalized by glycerol expression levels at the same condition. Statistical significance is represented as asterisks, considering p-value as at least < 0.05 (* < 0.05 < ** < 0.005 < *** < 0.0001 < ****).

The Ca^2+^-CRZ1-independent but synergistic mechanism was also observed when we analyzed the expression of some β-glucosidases coding genes after SCB and cellulose induction (Figure 4A-G, **Additional file 6: Figure S1 A-F**). Regarding β-glucosidases regulation, such mechanism of cooperation was so remarkable that only one of the studied genes (*cebl3b*) was modulated by CRZ1 alone (without Ca^2+^) (Figure 4D). Interestingly, in opposite to the main cellulases of *T. reesei* (*cel6a* and *cel7a*), β-glucosidases gene control tends to respond more effectively to cellulose stimuli, highlighting the role of these enzymes on plant cell wall deconstruction. It is also worth to notice that β-glucosidases coding genes were the class of genes which we identified the major number of putative CRZ1 binding sites in our *in silico* analysis, providing us an hypothesis of why these genes are more responsible to cellulose than other known strongly expressed genes (cellobiohydrolases).

**Figure 4.**
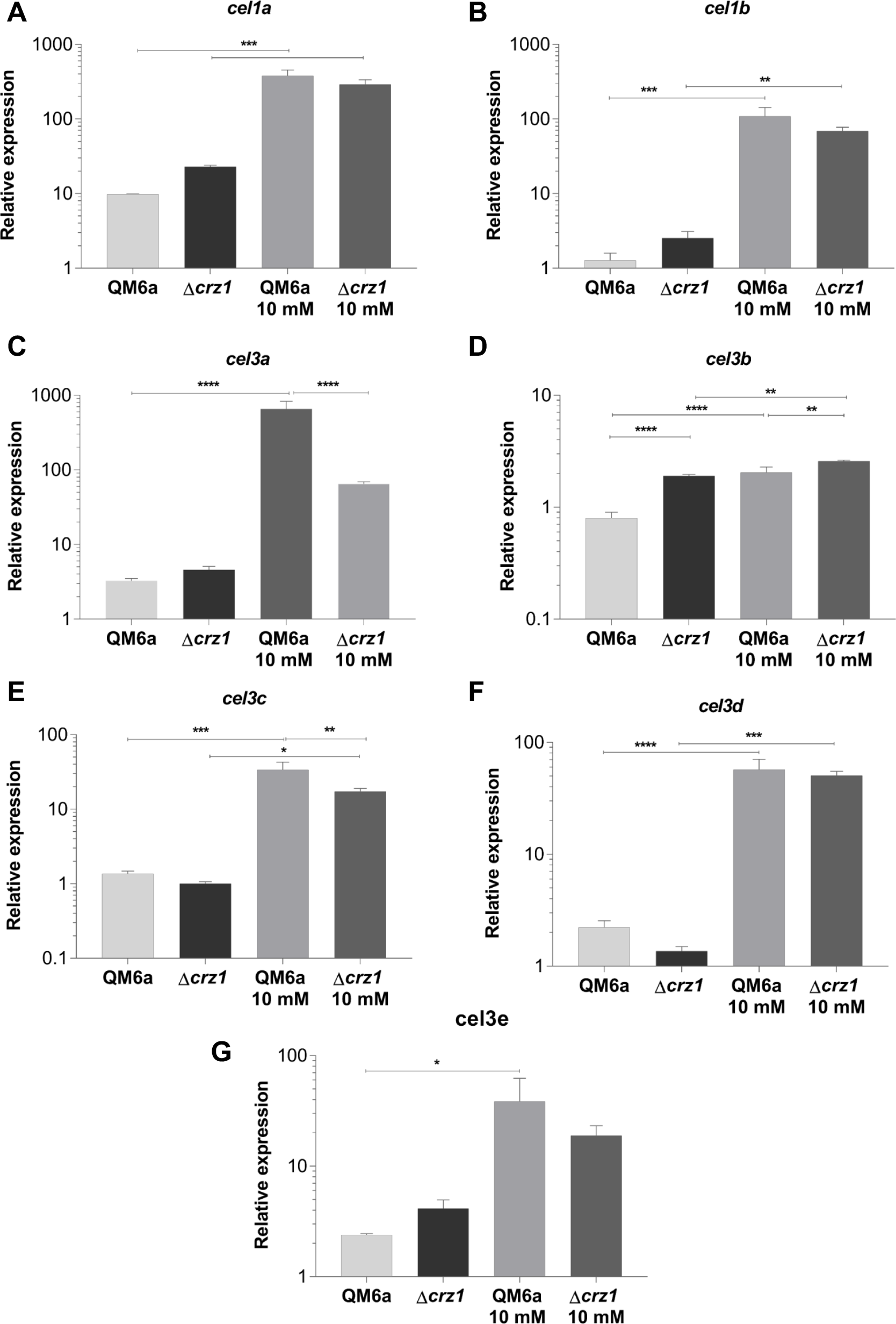
**A-G** – qRT-PCR results for differential expression analysis of the β-glucosidases genes of *T. reesei* in the QM6a and *Δcrz1* strains after 8 h of growing in SCB supplemented or not with 10 mM Ca^2+^. Expression values are represented as log_10_ means of three biological replicates with standard deviation normalized by glycerol expression levels at the same condition. Statistical significance is represented as asterisks, considering p-value as at least < 0.05 (* < 0.05 < ** < 0.005 < *** < 0.0001 < ****).

The most heterogeneous effect on holocellulases expression was noticed for endoglucanase coding genes expressed in SCB, as shown by Figure 5A-D. In this condition, we observed a Ca^2+^-CRZ1-independent decrease on endoglucanases expression (Figure 5A), a synergic Ca^2+^-CRZ1-independent effect which resulted in a increase on expression (Figure 5B,D) and finally, a positive modulation exclusively mediated by CRZ1 which loses its potential in a Ca^2+^-synergic mechanism (Figure 5C). Interestingly, after cellulose induction, the Ca^2+^-CRZ1-independent but synergistic mechanism was capable to potentialize the expression of only *cel7b* gene (**Additional File 7: Figure S1 B**).

**Figure 5.**
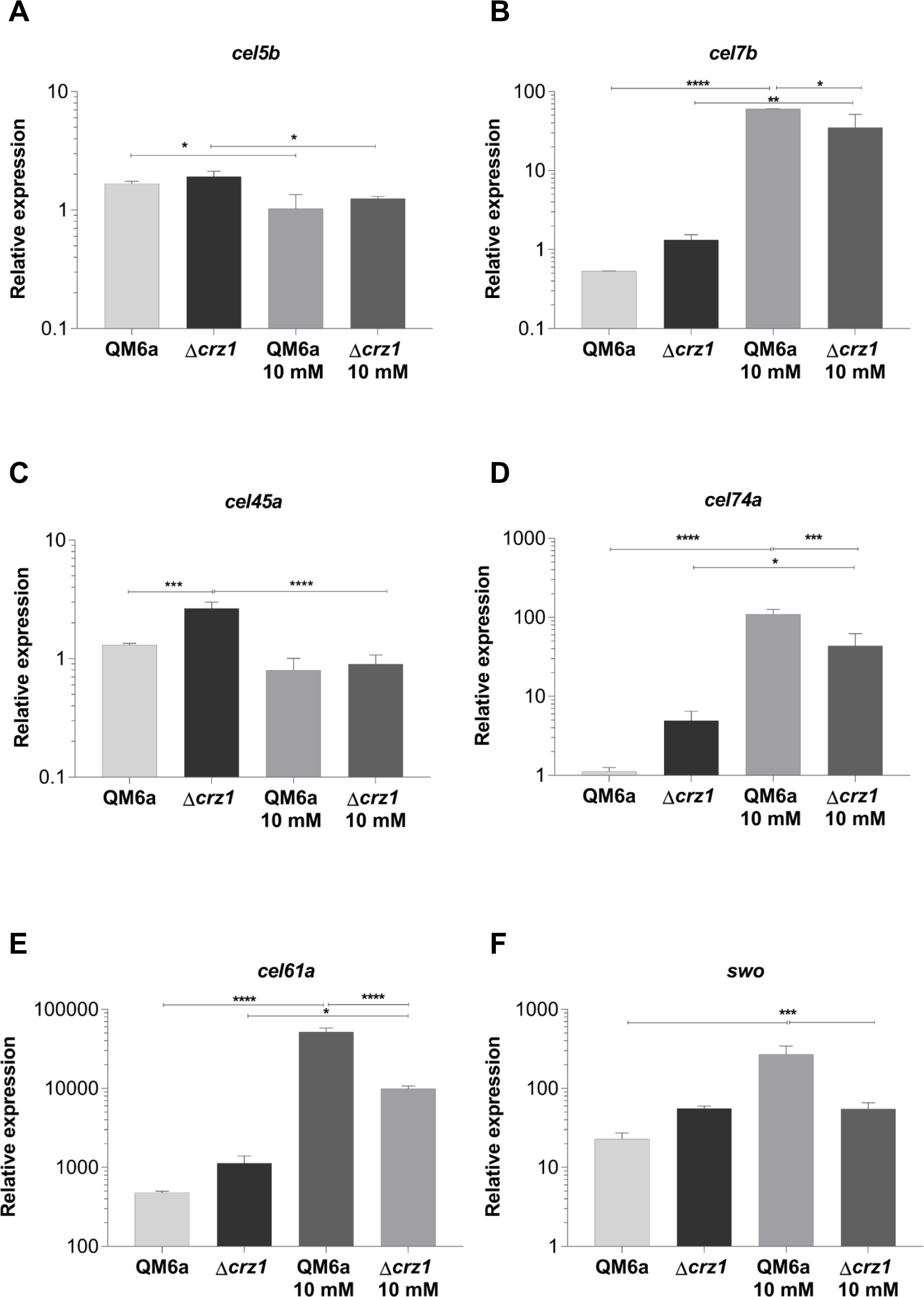
Analysis of differential expression of the endoglucanases (**A-D**), LPMO CEL61A (**E**) and swollenin (**F**) from QM6a and *Δcrz1 T. reesei* in the presence of SCB and SCB supplemented with 10 mM Ca^2+^ after 8h of induction. Expression values are represented as log_10_ means of three biological replicates with standard deviation normalized by glycerol expression levels at the same condition. Statistical significance is represented as asterisks, considering p-value as at least < 0.05 (* < 0.05 < ** < 0.005 < *** < 0 0001 < ****)

The effect of Ca^2+^ and the deletion of *crz1* on accessory proteins expression was also investigated in our study, as shown for the lytic polysaccharide monooxygenases (LPMO) *cel61a* and *swo* (swollenin, an expansin-like protein) genes. In this way, we also observed the effect of a synergistic Ca^2+^-CRZ1-independent mechanism that may positively regulate the expression of these genes in both SCB (Figure 5E-F) and cellulose induction (**Additional File 7: Figure S1 E-F**), highlighting the role of both CRZ1 and Ca^2+^ on the regulation of accessory proteins required for lignocellulose deconstruction.

### Calcium and sugar transporters are regulated by CRZl in the presence of Ca^2+^

Once the role of Ca^2+^ is clearly related to the CRZ1 regulatory role in *T. reesei* [26], we decided to investigate whether calcium transporters were target of modulation by CRZ1. After both SCB and cellulose growth, we observed that the deletion of *crz1* elicited a loss in transcription of these genes in the presence of SCB (Figure 6B-C) while it boosted the expression of the same genes after cellulose induction (**Additional File 8: Figure S1 B-D**). In spite of CRZ1-independent modulator effect, it is also worth to notice that Ca^2+^ synergistically to this TF was able to cause a decrease on the expression of such genes predominantly in SCB (Figure 8A-C) in comparison to cellulose (**Additional File 8: Figure S1 B**). Interestingly, our *in silico* analysis also evidenced the occurrence of putative CRZ1 binding sites in promoter regions of these genes, suggesting that calcium homeostasis is precisely regulated at genomic levels by CRZ1, mainly under stress conditions.

**Figure 6.**
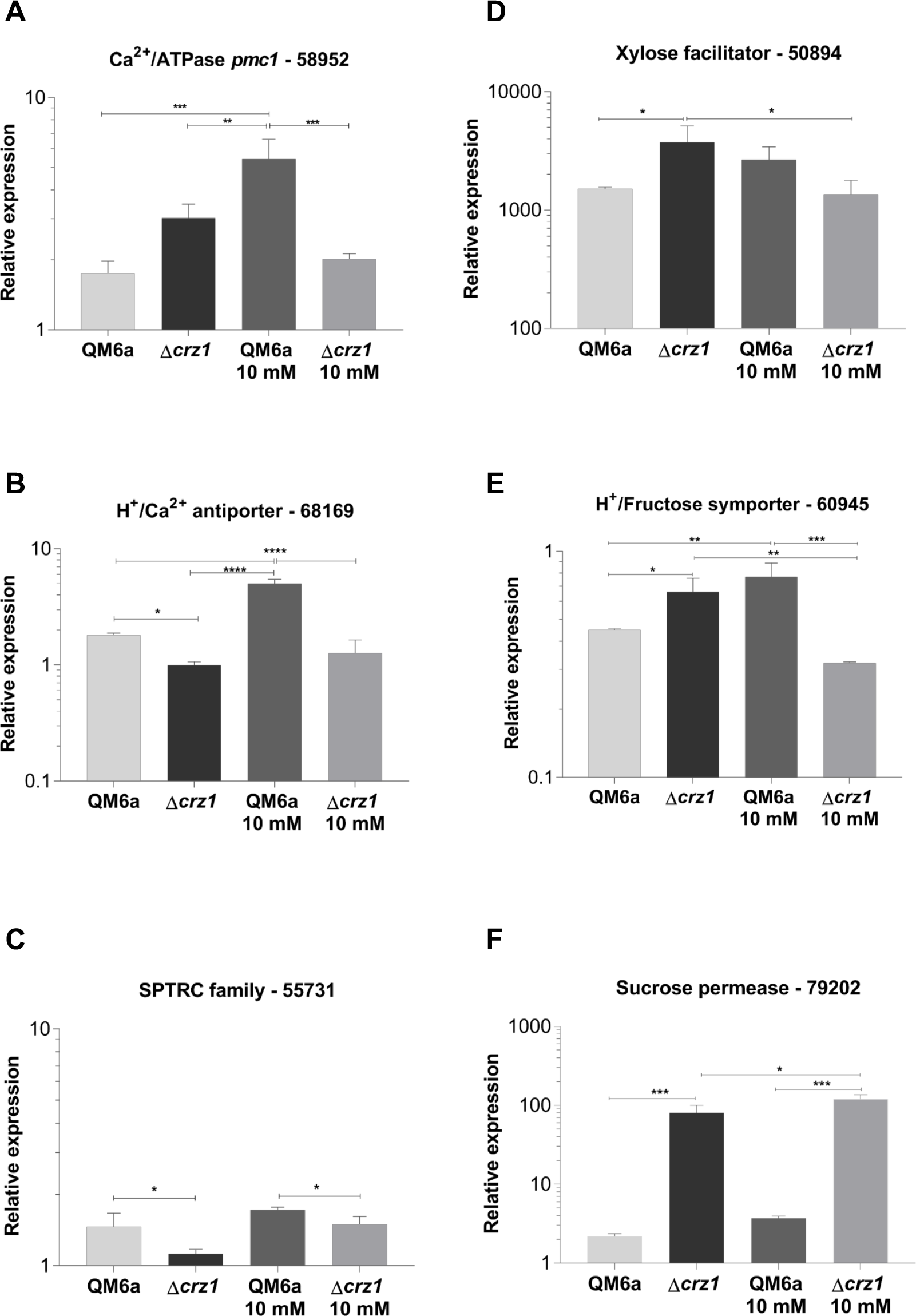
Ca^2+^ transporter genes (**A-C**) and sugar transporter (**D-F**) differential genic expression in the QM6a and *Δcrz1 T. reesei* strains after 8 h of growing in SCB and SCB supplemented with 10 mM Ca^2+^. Results are expressed as log_10_ means of three biological replicates with standard deviation normalized by glycerol expression levels at the same condition. Statistical significance is represented as asterisks, considering p-value as at least < 0.05 (* < 0.05 < ** < 0.005 < *** < 0.0001 < ****). Protein ID of the evaluated genes are supported at *T. reesei* genome database.

Finally, once we observed that carbon source could influence the transcription of holocellulases genes in *T. reesei* in Δ*crz1* strain, we studied the effects of *crz1* knockout on the expression of sugar transporter genes both after SCB and cellulose induction. Interestingly, our results showed that CRZ1 itself plays a regulatory role on the expression of these genes in the presence of complex substrates, since we detected a significant loss on their expression in SCB (Figure 6D-F). At the same conditions, we also observed a double effect on the synergic Ca^2+^ - CRZ1-independent mechanism. In this sense, Ca^2+^ allied to CRZ1, was able to significantly decrease the expression of *Trire_50894* and *Trire_60945* genes, while the opposite effect was achieved by the increase of *Trire_79202* gene expression. In contrast, concerning cellulose-induced differential regulation, we did not detect any differences on expression of the evaluated genes (**Additional file 9: Figure S1 C-E**), suggesting a remarkable role of CRZ1 in the regulation of genes involved in biomass transport.

### Effect of crz1 deletion on enzymatic activity of holocellulases

The analysis of the culture supernatants where the QM6a wild type and Δ*crz1* strains were grown showed that the activity of cellulases (β-glucosidases and endoglucanases) was always superior in the supernatants from Δ*crz1* strain in comparison to the QM6a strain, regardless the carbon source that we evaluated (Figure 7A-B, **Additional file 10: Figure S1 A-B**). In these assays, Ca^2+^ supplemenation was able to increase the enzymatic activity of all tested supernatants, mainly for endoglucanases, as we were not able to detect any enzymatic activity in the QM6a strain even in the presence of 10 mM Ca^2+^ for both carbon sources that we investigated.

**Figure 7.**
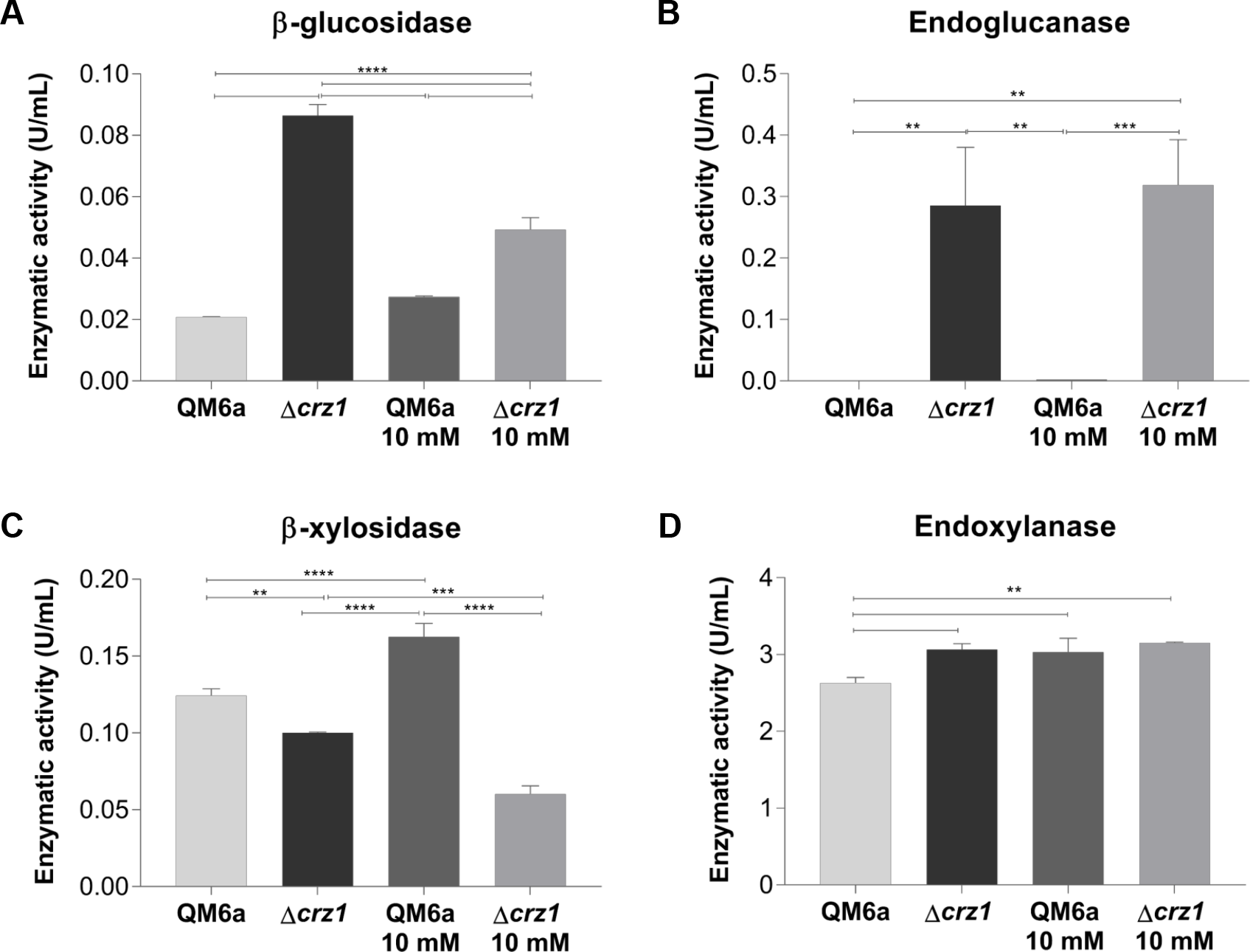
Enzymatic activity of β-glucosidase (**A**), endoglucanase (**B**), β-xylosidase (**C**) and endoxylanase (**D**) from QM6a and *Δcrzl T. reesei* strains supernatants after 8 h of growing in SCB and SCB supplemented with 10 mM Ca^2+^. Results are represented as absolute values in unities per milliliter and are representative of a mean of three biological replicates with standard deviation. Statistical significance is represented as asterisks, considering p-value as at least < 0.05 (* < 0.05 < ** < 0.005 < *** < 0.0001 < ****).

For xylanase activities (β-xylosidases and endoxylanases) in SCB and cellulose (Figure 7C-D, **Additional file 10: Figure S1 C-D**), we observed that in the Δ*crz1* supernatants the activities were higher compared to the QM6s strain except for the β-xylosidase in SCB (Figure 7C). Interestingly, the enzymatic activities of supernatants whose QM6a strains were grown after Ca^2+^ addition were higher in presence of SCB (Figure 7C-D) and presented a decrease in cellulose induction (**Additional file 10: Figure S1 C-D**). For instance, the *crz1* knockout resulted in supernatants with lower β-xylosidase activity in SCB presence (Figure 7C) and higher xylanases activities measures after cellulose induction (**Additional file 10: Figure S1 C-D**) when compared to the QM6a strain at the same condition.

## Discussion

In this study, we performed bioinformatics approaches to search for putative CRZ1 binding sites in promoter regions of holocellulases, Ca^2+^ and sugar transporter coding genes in *T. reesei* QM6a. Here we identified that differentially expressed holocellulases genes of this fungus [22] harbored sense and antisense CRZ1 putative binding sites at their ATG upstream sequences and it supports the evidence of the regulation played by this TF independently on carbon source availability. To be clear that CRZ1 was involved in gene regulation in the wild type strain of *T. reesei*, our study reports that holocellulases genes were directly or either indirectly modulated by this protein, corroborating the occurrences of putative CRZ1 binding sites sequences into *T. reesei* genome.

Concerning these findings, we observed that distinct classes of cellulases and xylanases were differentially expressed regardless the fungal growth condition. Although cellulose was not capable to modulate a vast number of genes independently on CRZ1 presence, SCB provided the major number of differentially expressed genes after its induction process (Figure 8A). This inducer capacity of SCB over the industrial strain RUT-C30 of *T. reesei* was already reported [54]. In their study, Borin *et al.* [54] found 24 biomass degrading enzymes in *T. reesei’s* RUT-C30 secretome after SCB exposure, showing that this carbon source is a potential activator of holocellulases production in this fungus. Despite RUT-C30 genome does not harness a *cre1* functional sequence, we verified here that for *T. reesei* QM6a, SCB was either the carbon source which preferentially elicited a higher holocellulases transcriptional response (Figure 8A).

**Figure 8.**
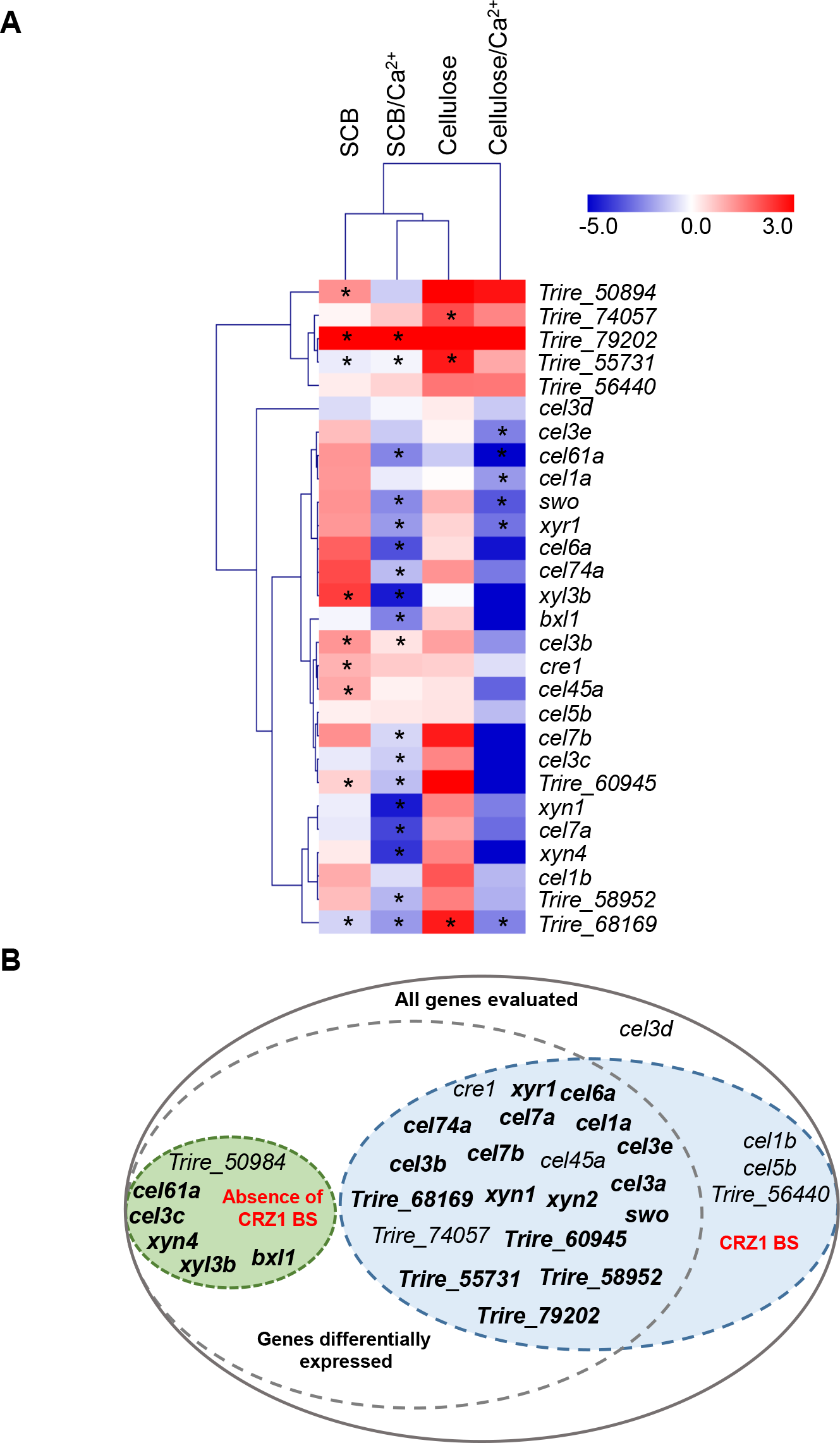
**A** – Heatmap of the differential expression profile of the holocellulases and calcium and transporter sugars after 8 h of induction in the presence or in the absence of 10 mM Ca^2+^. Values are representative of the expression obtained with mutant *crz1* strain related to the QM6a strain. Genes presenting representative statistical significance are show with asterisks. ID codes for calcium and sugar transporters are supported in *T. reesei* genome database. **B** – Representative scheme of genes differentially expressed in the absence of CRZ1 in *T. reesei* and its relation to the occurrence or absence of binding sites (BS) for CRZ1 at genomic level. Bold characters represents the genes whose expression was modulated by 10 mM Ca^2+^ addition.

Interestingly, our results also show that CRZ1-mediated regulation was not limited to lignocellulose degrading enzymes coding genes, but also intersected the regulatory network that coordinates holocellulases expression in the *T. reesei* QM6a. Therefore, we report that CRZ1 can modulate the transcript levels of relevant TFs such as XYR1 and CRE1 (inactive in RUT-C30 background), which in turn, can modulate cellulases and xylanases expression overall in this specie. This is specially interesting when we consider CRE1 repressor role on cellulases transcription, suggesting that deeper studies highlighting the interaction between CRZ1 and CRE1 may be a suitable strategy to engineer *T. reesei* as an industrial chassis for cellulases production.

Taking these facts in advantage, our data suggest a mechanism of synergy between the regulatory role of CRZ1 and the potential effect caused by Ca^2+^ upon the expression of holocellulases genes. In summary, we observed a major number of differentially expressed genes in presence of SCB supplemented with Ca^2+^ in comparison to cellulose exposure at the same Ca^2+^ concentration (Figure 8A), endorsing the previously reported effect of carbon source on modulation of *T. reesei* transcriptional profile [20]. In addition, these evidences corroborate the suggested synergic mechanism between CRZ1 and Ca^2+^, since we detected higher transcripts levels in all conditions supplemented with Ca^2+^ regardless the carbon source availability.

Ca^2+^ plays a regulatory role on several mechanisms in *T. reesei*, e.g. on protein secretion [55] and on events related to CRZ1-Calcineurin pathway [26]. In this sense, Chen *et al.* [26] reported the influence of 5 mM Ca^2+^ on the transcription of relevant industrial genes of *T. reesei* RUT-C30, such as *cbh1* and *xyr1*. In our study, 10 mM Ca^2+^ was able to potentialize the expression of *xyr1* at the first 8 hours of induction. This result is in contrast to previous findings reported by Chen *et al.* [26], whose study showed the first 6 hours of cellulose exposure as inefficient to increase expression of *xyrl* in the industrial strain of *T. reesei*. In conclusion, here we suggest that a higher Ca^2+^ concentration could elicit a faster gene activation in the presence of cellulose. Interestingly, we also observed a massive loss in transcription of *xyr1* in our Δ*crz1* strain in the presence of Ca^2+^, in accordance with the results obtained by Chen *et al.* [26] when authors investigated this effect in RUT-C30 strain, suggesting that CRZ1 is required for the activation of this relevant TF in *T. reesei*.

In the face of the pioneer contributions provided by Chen *et al.* [26], this study was performed using the RUT-C30 industrial strain of *T. reesei*, which does not harness CRE1, the master regulator of carbon catabolite repression in this fungus. Such genomic background could provide us an incomplete knowledge of the fully transcriptional CRZ1 pattern in these cells, mainly because in such transcriptional networks indirect regulation mechanisms may occur. For this reason, understand the mechanisms which control transcription in *T. reesei* QM6a could significantly contributes to the engineering holocellulases secretion in *T. reesei*.

Ca^2+^/CRZ1-Calcineurin/Calmodulin is one of the best studied signaling pathways which provides protection and sensibility for fungal cells against stress conditions. This signaling pathway is already reported to confer Ca^2+^ and cell wall stress tolerance in *Aspergillus* [56], pH responsiveness in *S. cerevisiae* [57] and it is either related to *C. neoformans* viability [29,33]. In addition, it is also reported that Ca^2+^ supplementation may influence distinct regulatory mechanisms that control holocellulases expression in *T. reesei* [26]. Although this signaling pathway are well characterized among fungal species, the effect of carbon source availability, Ca^2+^ concentration and CRZ1 dependence mechanisms together are still not completely clear in the field of cellulases regulation for the wild type strain of *T. reesei*.

In this sense, our data emphasize that Ca^2+^ is crucial for the transcription of such genes and this bivalent ion significantly contributed to the increase of transcript levels analyzed in our work through stimuli on Calmodulin/Calcineurin pathway. Furthermore, we were able to observe that Ca^2+^ seems to be critical for the expression of many holocellulases, Ca^2+^ and sugar transporter coding genes more directly than the carbon source whose fungal growth was previously established. Albeit the effect of SCB on enzymes expression, Ca^2+^ and sugar transporter proteins had already been reported in a *T. reesei* RUT-C30 gene co-expression network study by Borin *et al.* [5], here we firstly described the seminal importance of Ca^2+^ on the expression of sugar transporters in a CRZ1-dependent context in *T. reesei* QM6a strain, hypothesizing that under stress conditions *T. reesei* QM6a may increase the transcription of sugar transporters to keep metabolic homeostasis through Ca^2+^-CRZ1/Calcineurin pathway.

Finally, the effect of sense or antisense CRZ1 putative binding sites in promoter regions is one of the main evidences that support regulation QM6a strain of *T. reesei*. The effect of arrangements and repetitions of TFs binding sites in *T. reesei* was already reported by Kiesenhofer *et al.* [58] as an important parameter for engineering transcriptional regulation in this fungus. As observed in our work, through an *in silico* analysis, from the total set of differentially expressed genes, only 6 of them did not harness CRZ1 putative binding sites and only one of these was not modulated by Ca^2+^ (Figure 8B). These evidences strongly suggest that the synergic mechanism played by CRZ1 and Ca^2+^ is required for holocellulases and transporter coding genes expression in *T. reesei* QM6a even not in the presence of stress conditions. In addition, our data also assert that CRZ1 activation is pivotal for gene regulation, but Ca^2+^ is determinant for an increasingly modulation (positive or negative) on gene expression levels, claiming at the necessity of deeper studies on indirect holocellulases regulation in *T. reesei*.

## Conclusions

In this study, we highlight the major perspectives of bioinformatic approaches to extend our knowledge and expertise to engineer fungi for industrial applications. In summary, we describe the construction of a *T. reesei crz1* knockout strain through bioinformatics prediction and validation of the regulatory role of CRZ1 protein on transcriptional regulation of *T. reesei*. Allied to Ca^2+^, an important modulator of the regulatory effects of CRZ1, holocellulases and calcium/sugar transporter genes can be positively or negatively modulated in a crz1-synergic mechanism, as well as the enzymatic activity of such enzymes may direct or indirectly be affected by CRZ1 regulatory role. These phenomena are suitable attributes to biotechnological industry applications, highlighting fungi engineering as a fundamental platform to enzyme production at large scale.

