## Additional file for "CRZ1 regulator and calcium cooperatively modulate holocellulases gene expression in *Trichoderma reesei* QM6a"

**Additional file 1: Table S1.** Primers used in this study for the construction of the *crz1* deletion cassete.

| PRIMER | SEQUENCE |
|---|---|
| 5' Pcrz1 | CCAGTCCTGTTTCGCCATGTACGC |
| 3' Pcrz1-pyr4 | CCCAGACAAGACAAGGCAAGGAGGATGTGTCAAATAGGGCTGG |
| 5' pcrz1 -pyr4 | CCAGCCCTATTTGACACATCCTCCTTGCCTTGTCTTGTCTGGGTTCCT |
| 3' Pyr4 - Tcrz1 | TGCGACAACCCATCTAATCGGTACGGTTGATTGTTGCCGTCCGTTTC |
| 5' pyr4-Tcrz1 | GAAACGGACGGCAACAATCAACCGTACCGATTAGATGGGTTGTCGCA |
| 3' Tcrz1 | GCCATCCACCCCGAACTTCACAA |
| 5' Pcrz1 -pRS426 | GTAACGCCAGGGTTTTCCCAGTCACGACGCCAGTCCTGTTTCGCCATGTACGC |
| 3' Tcrz1 - pRS426 | GCGGATAACAATTTCACACAGGAAACAGCGCCATCCACCCCGAACTTCACAA |
| 5' Pcrz1 check1 | ATCCATACTACGGACCTTGTGCG |
| 3' pyr4 check1 | GTGTACTGCAGCTCGACGGTGT |
| 5' pyr4 check 2 | ATGCCTTTATCCACATGACGCCCG |
| 3' Tcrz1 check 2 | GCAATCAAAAGTCGCAGAAGACCG |
| PF crz1 qRT-PCR | CCCAAGAGATTCACCAGAGC |
| PR crz1 qRT-PCR | TTTCCTGTCATGCTGTCGAG |
| 5' crz1 ORF | TAAACCAATCCCACTCGCCG |
| 3' crz1 ORF | GCTCTGGTGAATCTCTTGGGAC |

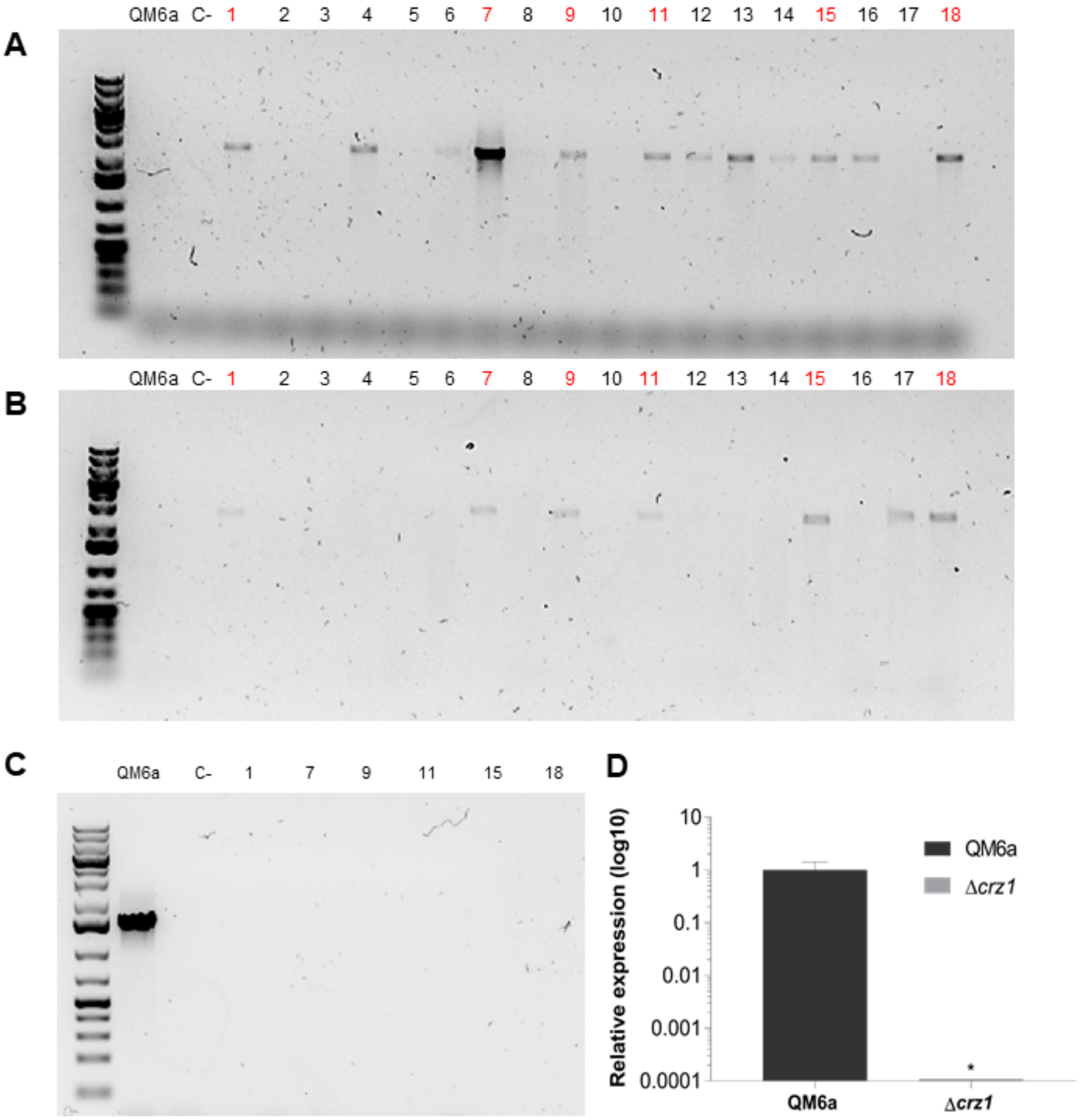


**Additional file 2: Figure S1.** Molecular procedures to confirm *crz1* ORF deletion from *T. reesei* QM6a genome. Numbers in lanes represent the numeric distribution of transformants obtained with the recombination. **A-** PCR whose amplicon of 2.6 kb comprises the annealing of a promoter external region primer and *pyr4* selectable marker. **B** – PCR to confirm the correct integration in *T. reesei* genome whose amplicon of 2.5 kb comprises the amplification of *pyr4* region with an external terminator sequence beyond the cassette of integration. **C** – Null amplification of the *crz1* ORF using a conventional PCR in the positive transformants previously obtained (**A-B**). **D-** Quantitative PCR of the *crz1* in the mutant strain obtained with homologous recombination.

**Additional file 3: Table S1.** Primers used in this study to quantify the differential expression levels of the holocellulases and calcium and sugar transporter genes in the *T. reesei Δcrz1* strain that we obtained.

*Holocellulases and swollenin primers*

| Name | Protein ID | Sequence |
|---|---|---|
| *cel6a* R | 72567 | TGT TCC ACC CGT TGT AGT TG |
| *cel6a* F | | ACA AGA ATG CAT CGT CTC CG |
| *cel3a* R | 76672 | TAG CTG AGA TCT CGT CGT C |
| *cel3a* F | | CTG TAC ATC ACC TAC CCA TC |
| *cel7a* R | 123989 | GGT AGC CTT CTT GAC TGA GT |
| *cel7a* F | | CCG AGC TTG GTA GTT ACT CTG |
| *cel1a* R | 120749 | AAT CAG CTC GTC AAA CAG CG |
| *cel1a* F | | TTT GCC TGG TCG CTC ATG |
| *cel61a* R | 73643 | ACC GCT GCC ACC ACA CTG |
| *cel61a* F | | GCG CCA CTG TTC CTG GAG |
| *cel45a* R | 49976 | TGG TCC AGA ATG CAC TCG |
| *cel45a* F | | CAG CGA CGT CTA CAT TGG |
| *cel7b* R | 122081 | AGG TCT TGG AGG TGT CAA CG |
| *cel7b* F | | CCC TCA ACA CTA GCC ACC AG |
| *cel1b* R | 22197 | TCC AAG TGC GAG TCA AAG TAG |
| *cel1b* F | | CCA TCT ACA TCA CCG AGA ACG |
| *cel3b* R | 121735 | ATG TGG AGG TTG GAG AAC TTG |
| *cel3b* F | | CCA GGA TAA CTT CAA CGA GGG |
| *cel3c* R | 82227 | ATC CCA ACC CCA TTC CTT TC |
| *cel3c* F | | GCG TAC AAT GGC ATC AAT GG |
| *cel3e* R | 76227 | TCG TGA GTC CAA AGT GAA CAG |
| *cel3e* F | | ATG TCT GGA AGT GAG GTT GC |
| *cel74a* R | 49081 | TGA TGT CTT TCC AAG TTC CCC |
| *cel74a* F | | GCC TTG TAT CTG ACC TAT TCC G |
| *xyn1* R | 74223 | TCT GTC TTT TGG GCT TGG AG |
| *xyn1* F | | GGC CAA ATT ATC GTC AAC TGT C |
| *xyn2* R | 123818 | TCT GCA CAG TAA CAG TTC CG |
| *xyn2* F | | TGT CAA CGA GCC TTC CAT C |
| *xyn4* R | 111489 | GAC AGG TTG GCA AAA TGG TG |
| *xyn4* F | | TGT CAG CAA TTC GGG TCT TC |
| *cel3c* F | 82227 | GCG TAC AAT GGC ATC AAT GG |
| *cel3c* R | | ATC CCA ACC CCA TTC CTT TC |
| *bxl1* F | 121127 | CAA GTC TGG AAT GAG GCT CTG |
| *bxl1*R | | TGA TGT CGG CAA TCT GGT G |
| *bxl3* F | 58450 | TCA ATG TTC CTC TCC ATG CC |
| *bxl3*R | | GGT TGG AGA TGG AGT AGA TGT TG |
| *swo* F | 123992 | CCA AAC TAT ACG AGT AGC C |
| *swo* R | | GAG TGA ATG TCT TGA TGG |
| *Actin* F | 44504 | TGA GAG CGG TGG TAT CCA CG |
| *Actin* R | | GGT ACC ACC AGA CAT GAC AAT GTT |

*Transcription factor primers*

| Name | Protein ID | Sequence |
|---|---|---|
| *xyr1* F | 122208 | CAA TCC TCT CCG TCG CTA TTC |
| *xyr1 R* | | CTG TTG CCG AAT GTG TTG AC |
| *cre1* F | 120117 | CTC CTA CTC GTC CTT TGT CAT G |
| *cre1* R | | GCA AGC ATC GTA ATG TCG TTG |
| *crz1* F | 36391 | CCC AAG AGA TTC ACC AGA GC |
| *crz1* R | | TTT CCT GTC ATG CTG TCG AG |

*Calcium and sugar transporter primers*

| Name | Protein ID | Sequence |
|---|---|---|
| Trire_58952 F | 58952 | CTC TCC AAA TGC TAC CCT TAC C |
| Trire_58952 R | | CCG ACT ACA TCC GCC ATG |
| Trire_55731 F | 55731 | ACG AAT CTA CCC TTT GCT GG |
| Trire_55731 R | | CGT AGT CGG TTG TCC TTC ATC |
| Trire_56440 F | 56440 | TGA GTG AAT GGA TAG ATG CCG |
| Trire_56440 F | | AGA GGG TGC CAA TGA GAA TC |
| Trire_74057 F | 74057 | CGT CAT GGG TGT TCT ATA CTC G |
| Trire_74057 R | | AAT TGC TCC CAC TCT TCC AC |
| Trire_68169 F | 68169 | TCT GGT TCA ACT TCC TCG C |
| Trire_68169 R | | CAC TCA TAG GAC CCA ACA CG |
| Trire_50894 F | 50894 | TCG TCT TTC GTC TCA TTG TCG |
| Trire_50894 R | | GAT TGC CTG TTT GTC TTC CG |
| Trire_79202 F | 79202 | GAG TAC CTG GTC AAC GTG AG |
| Trire_79202 R | | CAA ACC TAT GTG TTG GCT CAG |
| Trire_60945 F | 60945 | GTC TCT GGG CGT GTT ATT CTC |
| Trire_60945 R | | CTT CTC CAA GCG CAA TGT TG |

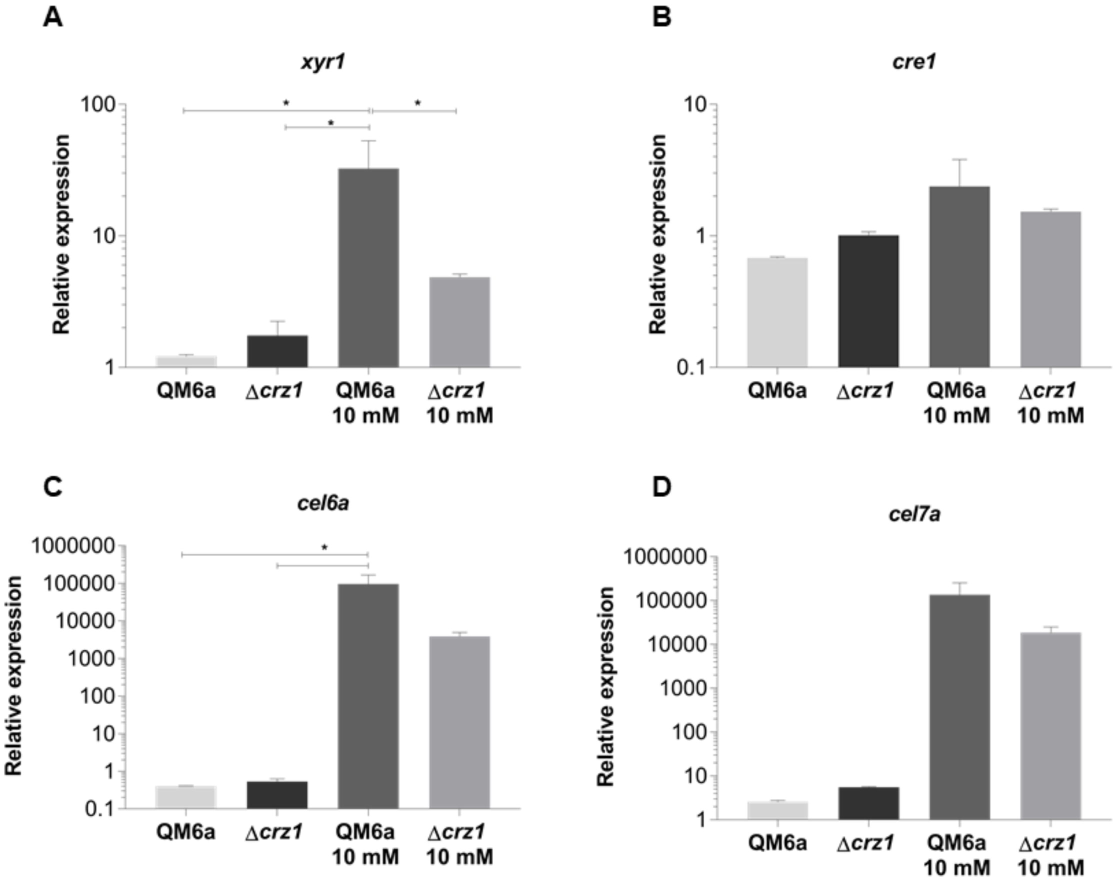


**Additional file 4: Figure S1** – Analysis of differential expression of the transcription factors XYR1 (**A**), CRE1 (**B**) and of cellobiohydrolases CEL6A (**C**) and CEL7a (**D**) from QM6a and *Δcrz1 T. reesei* strains after 8 h of growing in commercial cellulose (Avicel – Sigma Aldrich®) supplemented or not with 10 mM $Ca^{2+}$. Expression values are represented as $log_{10}$ means of three biological replicates with standard deviation normalized by glycerol expression levels at the same condition. Statistical significance is represented as asterisks, considering p-value as at least < 0.05 (* < 0.05 < ** < 0.005 < *** < 0.0001 < ****).

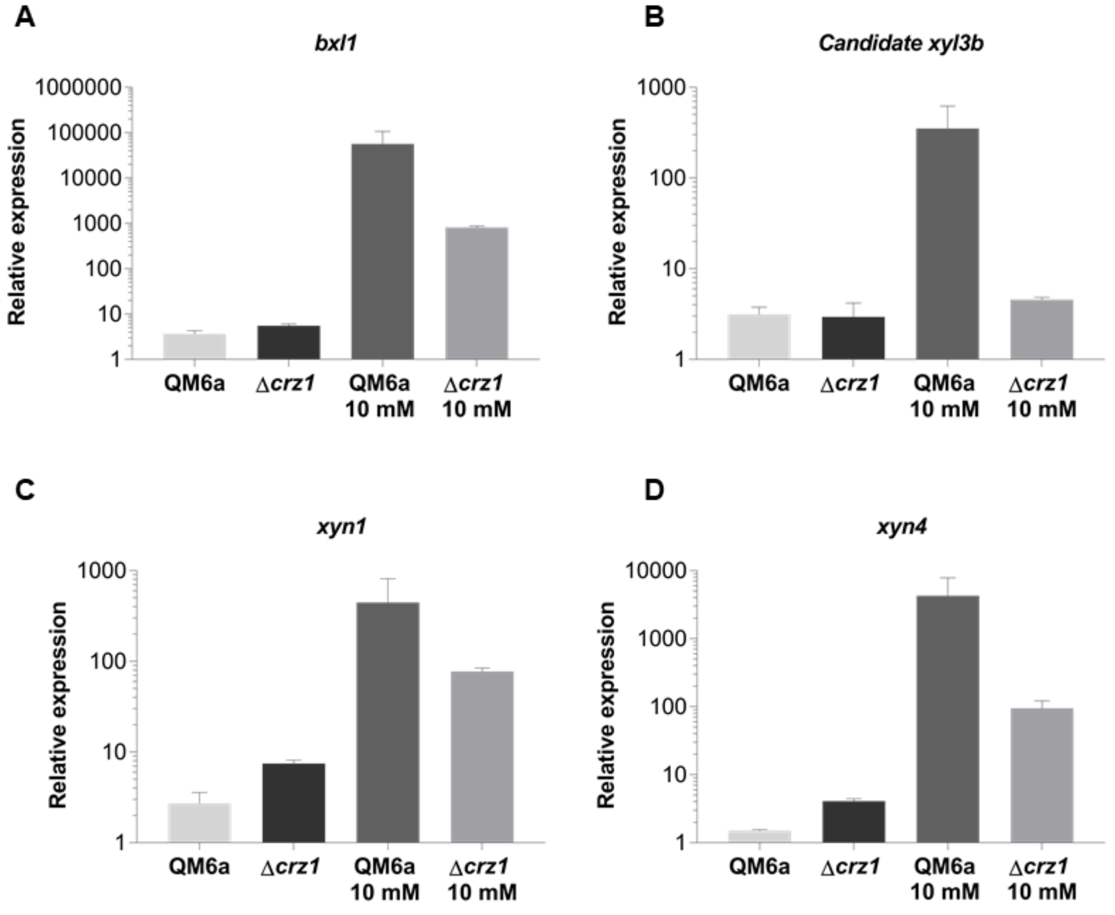


**Additional file 5: Figure S1 -** Relative expression of the β-xylosidase 1 (**A**) and the candidate *xyl3b* xylosidase (**B**) and of the endoxylanases *xyn1* (**C**) and *xyn4* genes (**D**) from QM6a and *Δcrz1 T. reesei* strains after 8 h of growing in commercial cellulose (Avicel – Sigma Aldrich®) supplemented or not with 10 mM $Ca^{2+}$. Expression values are represented as $log_{10}$ means of three biological replicates with standard deviation normalized by glycerol expression levels at the same condition. Statistical significance is represented as asterisks, considering p-value as at least < 0.05 (* < 0.05 < ** < 0.005 < *** < 0.0001 < ****).

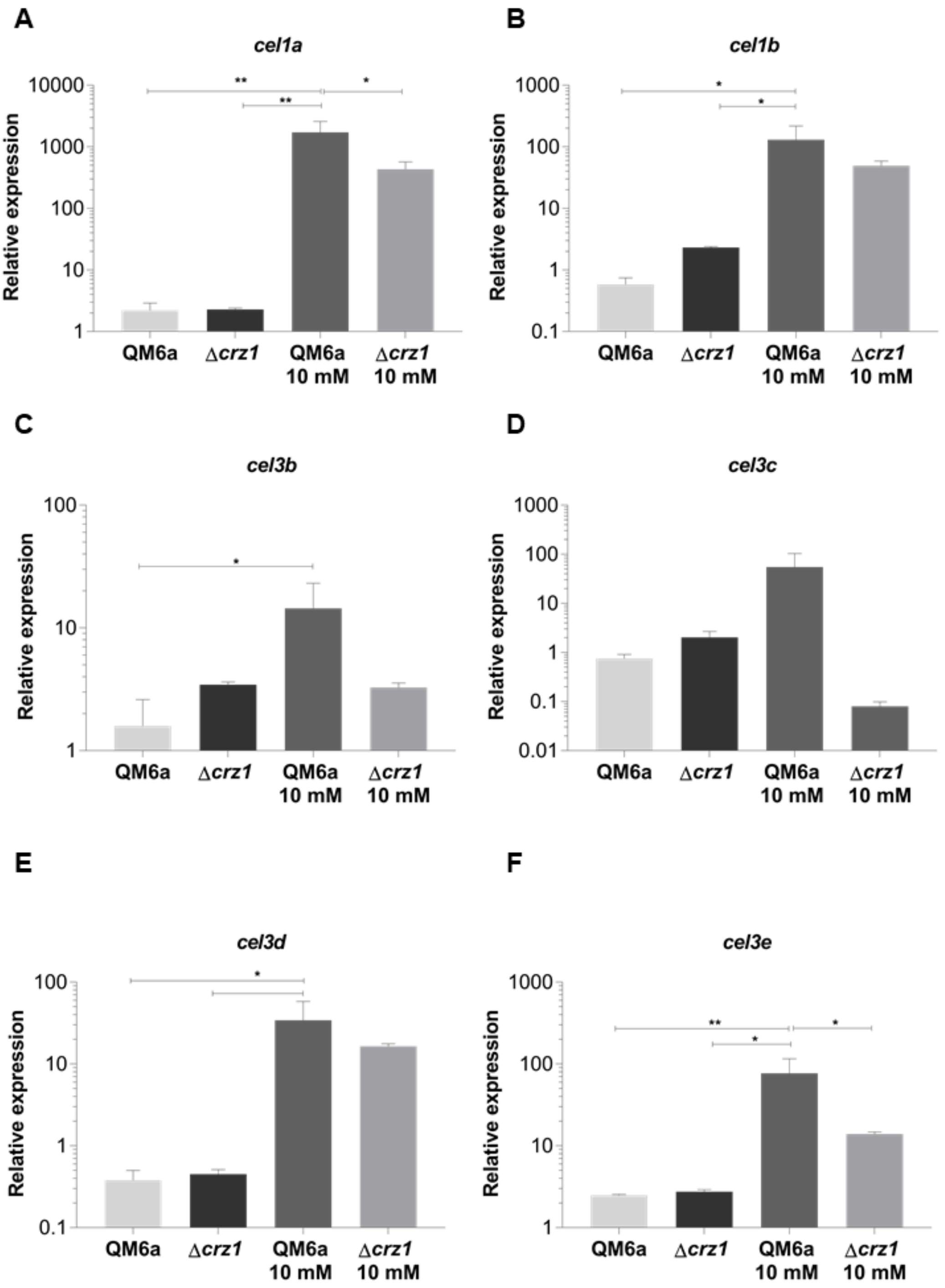


**Additional file 6: Figure S1 A-F -** Analysis of differential expression of the β-glucosidases genes of *T. reesei* in the QM6a and *Δcrz1* strains after 8 h of growing in commercial cellulose (Avicel – Sigma Aldrich®) supplemented or not with 10 mM $Ca^{2+}$. Expression values are represented as $log_{10}$ means of three biological replicates with standard deviation normalized by glycerol expression levels at the same condition. Statistical significance is represented as asterisks, considering p-value as at least < 0.05 (* < 0.05 < ** < 0.005 < *** < 0.0001 < ****).

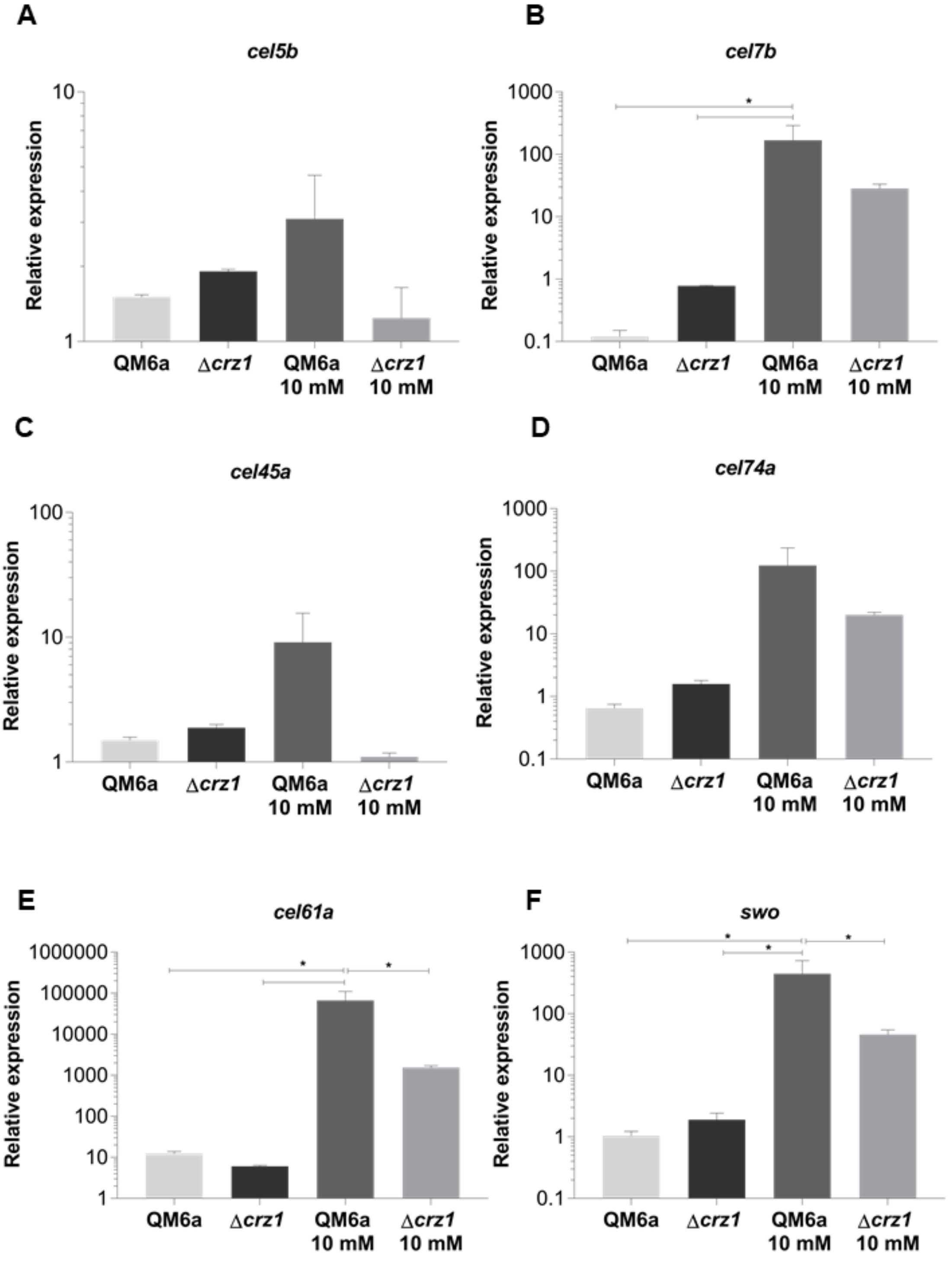


**Additional file 7: Figure S1** – Endoglucanases (**A-D**), LPMO CEL61A (**E**) and swollenin (**F**) differential expression from the QM6a and *Δcrz1* strains after 8 h of growing in commercial cellulose (Avicel – Sigma Aldrich®) supplemented or not with 10 mM $Ca^{2+}$. Expression values are represented as $log_{10}$ means of three biological replicates with standard deviation normalized by glycerol expression levels at the same condition. Statistical significance is represented as asterisks, considering p-value as at least < 0.05 (* < 0.05 < ** < 0.005 < *** < 0.0001 < ****).

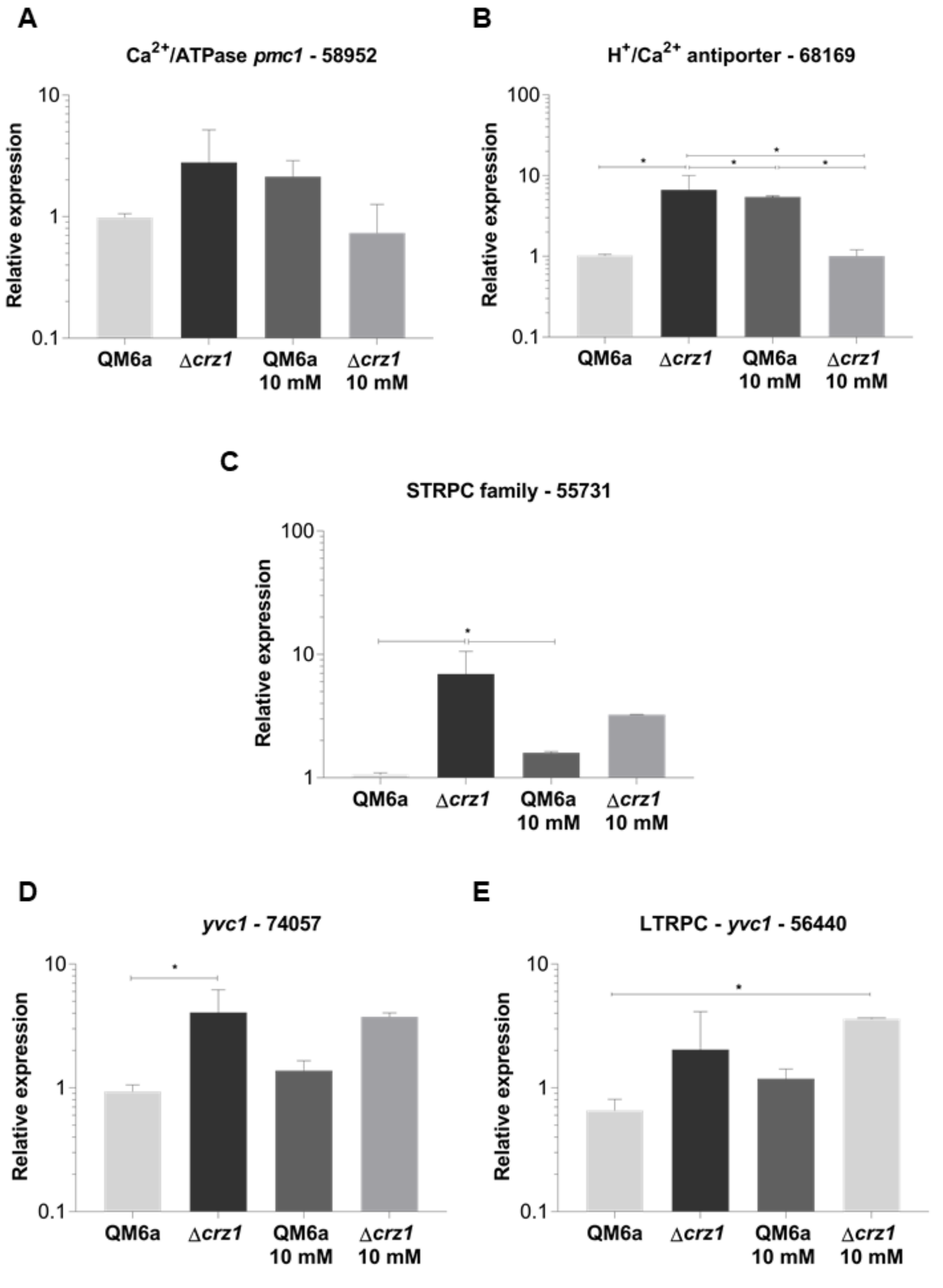


**Additional file 8: Figure S1. A-E** – qRT-PCR results for differential expression analysis of the Calcium-transporter genes differential expression in the QM6a and *Δcrz1 T. reesei* strains after 8 h of growing in commercial cellulose (Avicel – Sigma Aldrich®) supplemented or not with 10 mM $Ca^{2+}$. Expression values are represented as $\log_{10}$ means of three biological replicates with standard deviation normalized by glycerol expression levels at the same condition. Statistical significance is represented as asterisks, considering p-value as at least < 0.05 (* < 0.05 < ** < 0.005 < *** < 0.0001 < ****). Protein ID of the evaluated genes are available at *T. reesei* genome database (https://genome.jgi.doe.gov/pages/search-for-genes.jsf?organism=Trire2).

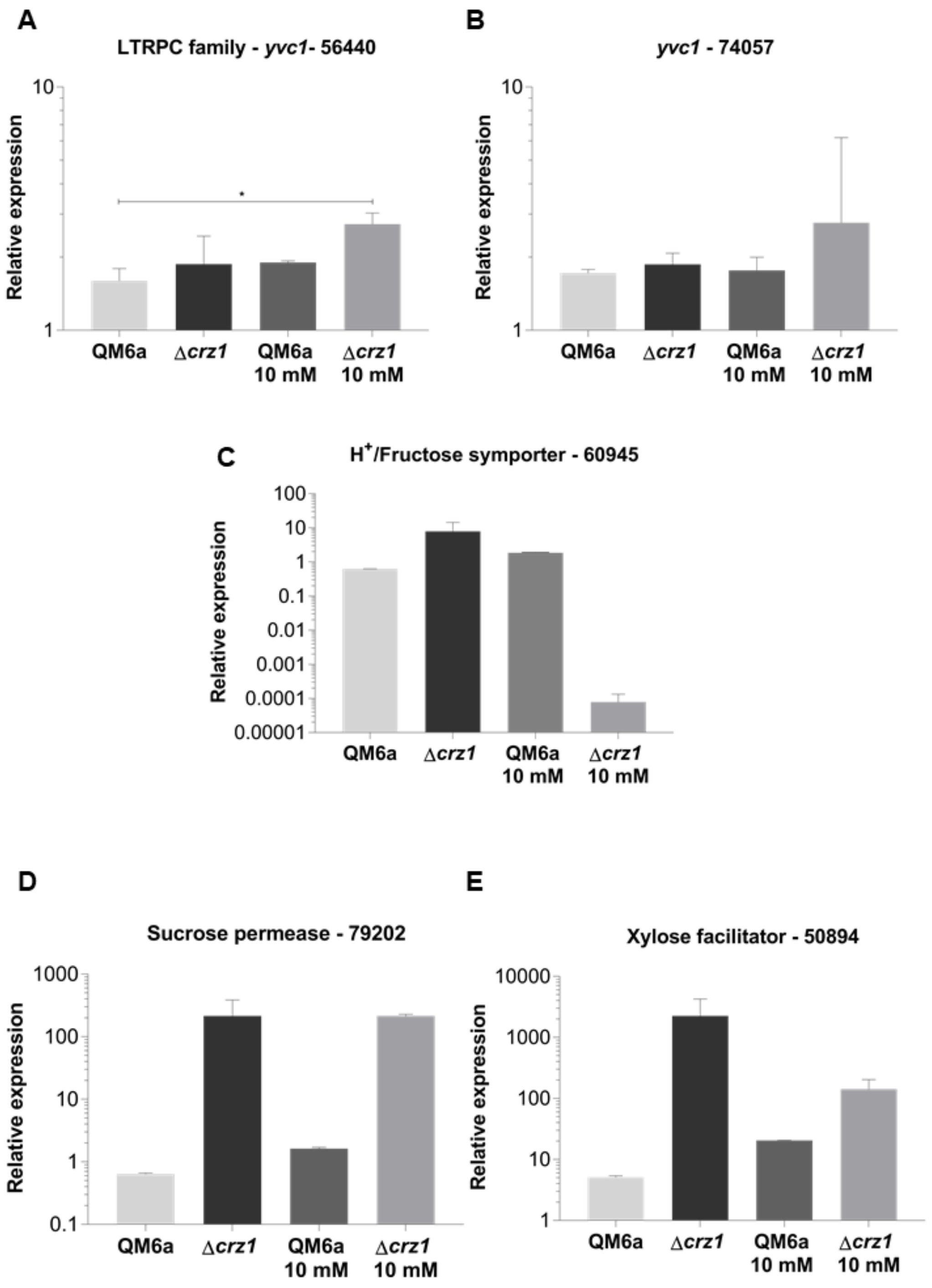


**Additional file 9: Figure S1 A-B** – qRT-PCR results for differential expression analysis of the Calcium-transporter genes differential expression in the QM6a and *Δcrz1 T. reesei* strains after 8 h of growing in SCB supplemented or not with 10 mM $Ca^{2+}$. **C-E -** Sugar transporter genes differential expression analysis in the QM6a and *Δcrz1 T. reesei* strains grown in commercial cellulose (Avicel – Sigma Aldrich®) supplemented or not with 10 mM $Ca^{2+}$ after 8 h of induction. Expression values are represented as $\log_{10}$ means of three biological replicates with standard deviation normalized by glycerol expression levels at the same condition. Statistical significance is represented as asterisks, considering p-value as at least < 0.05 (* < 0.05 < ** < 0.005 < *** < 0.0001 < ****). Protein ID of the evaluated genes are available at *T. reesei* genome database (https://genome.jgi.doe.gov/pages/search-for-genes.jsf?organism=Trire2).

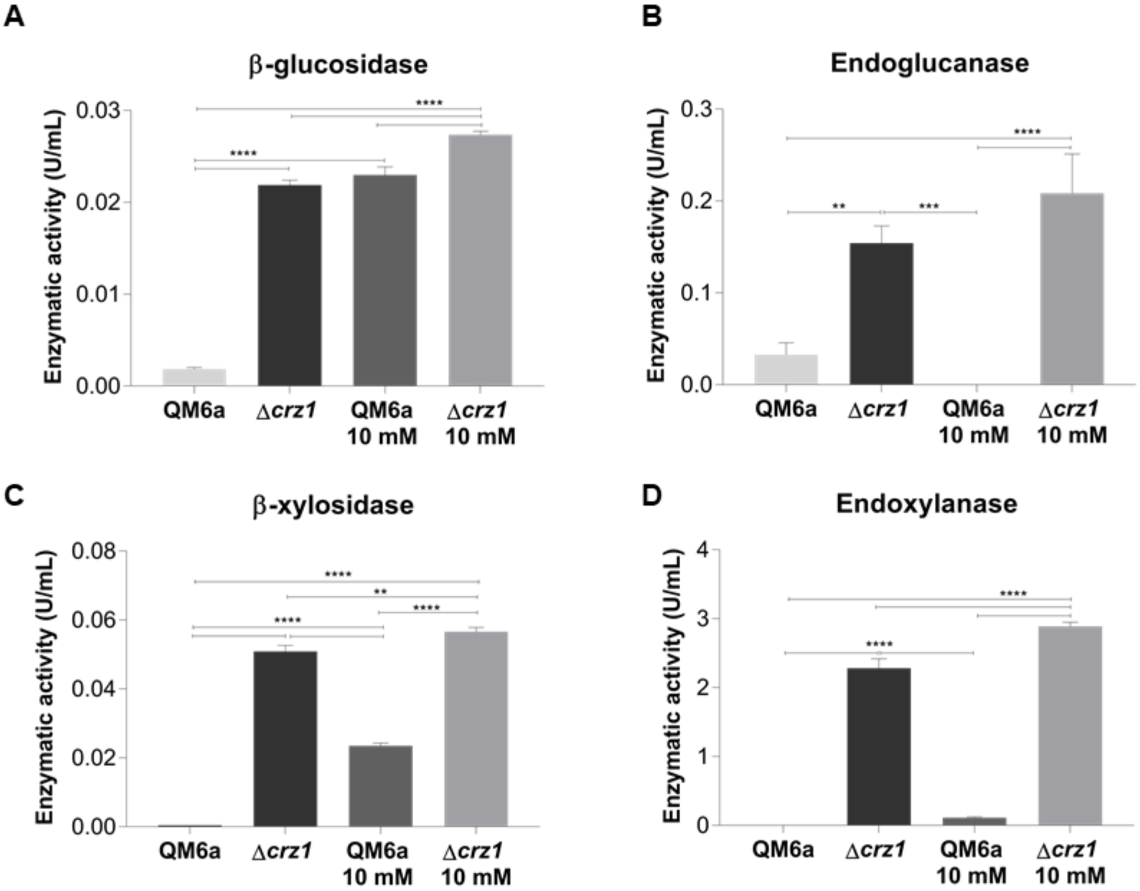


**Additional file 10: Figure S1 -** Enzymatic activity measurements for β-glucosidase (**A**), endoglucanase (**B**), β-xylosidase (**C**) and endoxylanase (**D**) from QM6a and *Δcrz1 T. reesei* strains supernatants after 8 h of induction with commercial cellulose (Avicel – Sigma Aldrich®) supplemented or not with 10 mM $Ca^{2+}$. Results are represented as absolute values in unities per milliliter and are representative of a mean of three biological replicates with standard deviation. Statistical significance is represented as asterisks, considering p-value as at least < 0.05 (* < 0.05 < ** < 0.005 < *** < 0.0001 < ****).
